# Mechanical cues stabilize a conserved morphometric state associated with radial glia-like competence across systems

**DOI:** 10.1101/2025.01.09.632137

**Authors:** J.P. Soriano-Esqué, C. Borau, R. Sortino, L.D. Garma, J.A. Ortega, J. Asín, S. Alcántara

## Abstract

Mechanical cues influence neural development, yet how tissue architecture is integrated into progenitor cell states remains poorly understood. Here, we show that defined microtopographies induce an early nuclear remodeling program associated with radial glia (RG)-like competence. Aligned microgrooves promote nuclear elongation, reduced Lamin A/C to B1 ratio, sustained β-catenin activity, and distinct patterns of nuclear calcium dynamics, all of which merge before peak Pax6 expression. Pharmacological inhibition of mechanosensitive calcium signaling abolishes RG-associated marker induction while preserving nuclear remodeling, indicating that calcium-dependent pathways are required for Pax6 induction but are dispensable for the establishment of the underlying morphometric state. To quantitatively describe these transitions, we developed an interpretable model based on nuclear geometry and local cell density that identifies morphometric states associated with RG-like competence across experimental conditions. Application of this framework to embryonic mouse and human cortex revealed analogous signatures in native RG populations. Together, these findings indicate that tissue architecture influences RG-like competence through conserved nuclear morphometric states across developmental contexts.

**HIGHLIGHTS:**

- Microtopography elicits coordinated cellular and nuclear remodeling linked to RG-like competence.
- Nuclear morphometric states precede peak expression of RG-associated markers.
- Calcium signaling is required for Pax6 induction but is dispensable for early structural remodeling.
- Conserved morphometric states associated with RG-like competence emerge across mouse, human, and organoid systems.

## INTRODUCTION

The adult mammalian central nervous system (CNS) exhibits a strikingly limited capacity for self-repair despite the persistence of neural stem cells (NSCs). During embryogenesis, radial glia (RG) constitutes the primary NSC population, generating neurons, glial cells, and intermediate progenitors (IPs)^1,2^. RG are abundant and highly proliferative during corticogenesis, supporting neurogenesis, cortical lamination and the emergence of complex cytoarchitecture. Shortly after birth, however, most RG differentiate into astrocytes^1–3^, and only a small subset persists as RG-like NSCs within restricted neurogenic niches^4,5^. Understanding how RG-associated states are established and maintained is therefore central to both developmental and regenerative neurobiology.

RG exhibit a characteristic bipolar morphology spanning the apical ventricular surface and the basal pia. These apical-basal anchorages generate mechanical tension critical for maintaining RG identity^6–8^. Disruption of these anisotropic architectural constraints produces distinct progenitor populations: partial loss of anchorage gives rise to outer and truncated RG (oRG and tRG), which retain polarity and expression markers such as Pax6, Sox2, and Nestin^3^. In contrast, complete loss of polarity produces IPs expressing TBR2/EOMES^9^, NG2^10^, or GSX2^11^. These observations suggest that RG-associated states are tightly linked to structural and mechanical inputs from the physiological niche. Consistent with this view, RG to astrocyte transitions are dynamic and context-dependent, with astrocytes typically identified by the expression of markers such as GFAP, S100β, and glutamine synthetase^3^. Importantly, these conversions are bidirectional^12,13^: following injury, astrocytes undergo marked changes in morphology, calcium dynamics, and transcriptional state^14–18^, and a subset reacquires RG-like properties^16,19–21^, indicating that astrocytes may act as a reservoir of latent stem cell potential. These transitions highlight the plasticity of RG-associated states and suggest that cell identity may be reversible under appropriate niche-derived mechanical and signaling cues.

Mechanical cues are increasingly recognized as important regulators of neural progenitor behavior. Mechanotransduction, the process by which physical forces are converted into biochemical signals, operates through cytoskeletal tension, nuclear deformation, mechanosensitive ion channels (MSCs), Ca²⁺ signaling, and chromatin reorganization^22–25^. Both astrocytes and neural progenitors, including RG, are highly mechanosensitive, and substrate mechanics profoundly influence their proliferation, differentiation, and functional state^26–29^. However, how these diverse inputs are integrated into coherent cell states remains unclear, particularly with respect to the coordination of nuclear architecture and calcium dynamics.

Engineered biomaterials provide a powerful framework to dissect how defined mechanical cues shape cell behavior^30–36^. We previously developed a non-biodegradable poly(methyl methacrylate) substrate with 2 µm linear grooves (ln2PMMA) that recapitulates key aspects of RG anisotropic microtopography^34,35^. Astrocytes cultured on this substrate acquired RG-like features, including bipolar morphology, expression of Pax6 and Sox2, and the ability to support neuronal migration^34,35^. In ln2PMMA, astrocyte-RG transitions are bidirectional and cell density dependent, with cells maturing into astrocytes upon reaching confluency and reverting to RG-like phenotypes after reseeding at the original cell density^35^.

These findings suggest that microtopography can stabilize RG-like states; however, the mechanistic links among anisotropic cues, nuclear remodeling, calcium signaling, and RG-associated marker induction remain poorly understood.

Here, we investigate how mechanically induced astrocyte to RG-like transitions are associated with changes in nuclear architecture, calcium signaling, and cellular morphology. We identify a conserved morphometric state linked to RG-like competence, characterized by coordinated changes in nuclear architecture, cellular morphology, and calcium dynamics. To quantitatively capture this state, we develop an interpretable morphometric statistical model based on nuclear geometry and local cell density. We further apply this framework to human cortical tissue, revealing conservation of RG-associated morphometric features across systems. Together, this work provides a framework for understanding how mechanical cues promote RG-like competence through coordinated changes in nuclear organization, cellular morphology, and calcium signaling.

## RESULTS

### Mechanical cues induce early nuclear remodeling preceding RG-associated marker induction

To investigate how mechanical cues contribute to the emergence of RG-like states, we analyzed Pax6, Nestin, and GFAP expression in astrocyte cultures grown on uncoated ln2PMMA, an RG-mimetic substrate with aligned microgrooves^34,35^, from 1 to 14 days *in vitro* (DIV), using uncoated glass as the control substrate (**Fig. 1A, B**). Pax6 expression, used as a readout of RG-associated marker induction, was similar between ln2PMMA and the control substrate at 1 DIV. By 3 DIV, however, ln2PMMA cultures showed a significant decrease in GFAP⁺ astrocytes together with an increase in Pax6⁺ cells, a difference that persisted at 7 DIV and converged again at 14 DIV (**Fig. 1C**). These dynamics are consistent with previous studies showing that mechanically induced RG-like states are reversible and sensitive to substrate geometry and cell confluence^34,35^. To determine whether changes in lineage marker expression induced by mechanical cues act through early nuclear remodeling, we quantified nuclear orientation and elongation (**Fig. 1A**). Across all time points, nuclei on ln2PMMA were elongated and aligned with the grooves, whereas nuclei on flat substrates remained rounded and randomly oriented (**Fig. 1D**). Notably, nuclear alignment and elongation were already established after 24 h of culture, when Pax6 frequencies remained indistinguishable between substrates, indicating that structural remodeling precedes detectable RG-associated marker induction. This temporal relationship is consistent with the idea that nuclear deformation is an early mechanosensitive response linked to subsequent RG-associated marker induction. These observations indicate that nuclear remodeling emerges before the induction of RG-associated markers, suggesting that mechanical cues initially establish a permissive morphometric state associated with RG-like competence.

**Figure 1.**
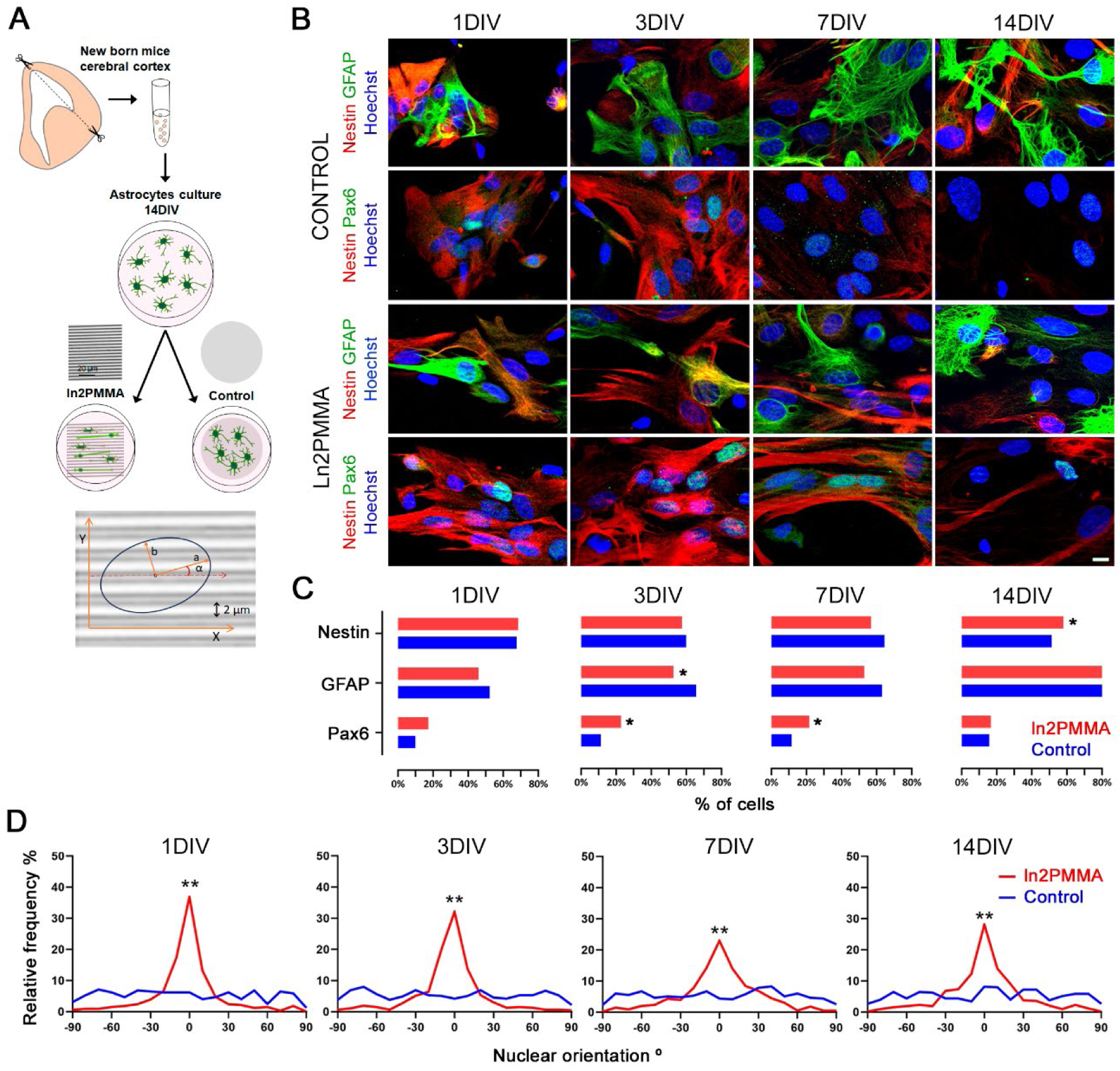
Glial cell composition and nuclear alignment on ln2PMMA and control substrates. **(A)** Schematic representation of the experimental design and the workflow used for nuclear morphology parameterization. **(B)** Representative images of glial cultures at 1, 3, 7, and 14 days *in vitro* (DIV) grown on control glass and ln2PMMA substrates. Cells were immunolabeled for Nestin (red) and GFAP (green), or for Pax6 (green). Nuclei were counterstained with Hoechst (blue). **(C)** Quantification of Nestin⁺, Pax6⁺, and GFAP⁺ cell populations on each substrate at the time points shown in **(B)**. **(D)** Relative frequency distributions of nuclear orientation at the indicated time points. Angles were converted from a 0-360° range to a –90° to 90° range, with 0°corresponding to alignment with the substrate topography. A total of 2,135 nuclei from control cultures and 1,720 nuclei from ln2PMMAcultures were analyzed. Frequencies of marker-defined cell populations were compared using a chi-square test, and nuclear orientation values were compared using the Mann-Whitney test. *p < 0.05; **p < 0.01. Scale bar = 10 µm.

We next asked whether nuclear shape remodeling was accompanied by changes in the nuclear lamina. Immunofluorescence quantification revealed significantly lower Lamin A/C to B1 ratios in ln2PMMA cultures at 1, 3, and 7 DIV (**Fig. 2A-B**), consistent with a more compliant lamina state typical of RG nuclei^37,38^. Because nuclear lamina composition modulates mechanosensitive transcriptional regulators^39,40^, we examined the localization of β-catenin and YAP/TAZ, as readouts of pathway activation. Nuclear β-catenin levels were higher at 1 DIV and decreased more rapidly in control cultures, whereas ln2PMMA cultures retained higher nuclear β-catenin levels over time (**Fig. 2C-D**). In contrast, no differences in YAP/TAZ localization were detected at the analyzed time points (**Fig. 2E-F**). Given the reported link between nuclear β-catenin and Pax6 regulation^41^, the sustained β-catenin nuclear retention in ln2PMMA cultures is consistent with this association. This observation supports a link between mechanically induced nuclear remodeling, sustained β-catenin activity, and the emergence of RG-associated states. However, as YAP/TAZ binds to β-catenin^42^, we cannot completely rule out its involvement, and a direct causal relationship remains to be established.

**Figure 2.**
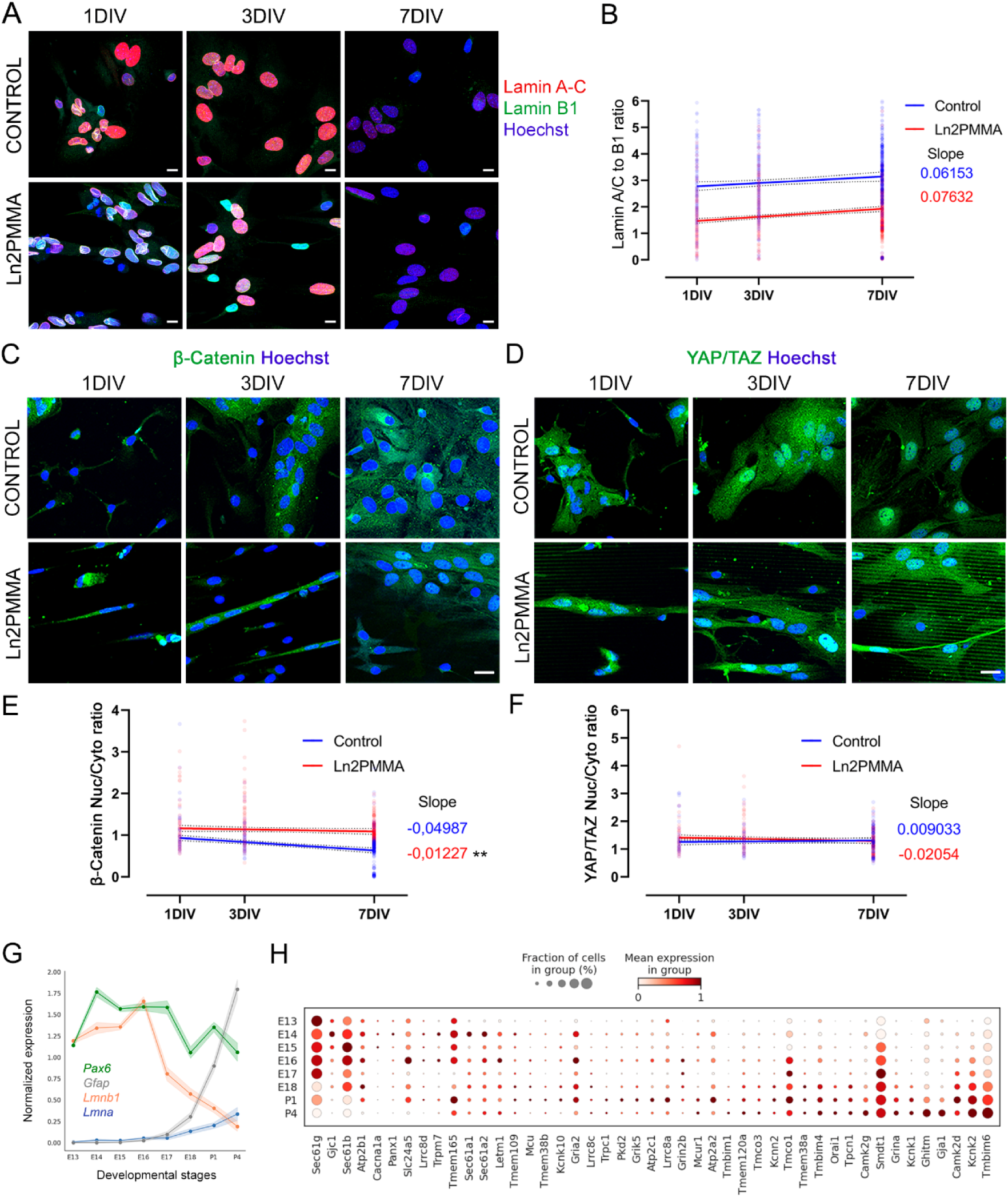
Changes in mechanotransduction effectors during RG to astrocyte state transitions. **(A)** Confocal images of glial cells cultured for 1-7 DIV on control and ln2PMMA substrates and immunostained for lamin A/C (red) and lamin B1 (green). Nuclei were counterstained with Hoechst 33342 (blue). **(B)** Linear regression analysis of nuclear lamin A/C to B1 intensity ratio in glial cells across time in culture (control n = 780; ln2PMMA n = 792). **(C)** Confocal images of β-catenin immunostaining (green). **(D)** Linear regression analysis of β-catenin nuclear to cytoplasmic (Nuc/Cyto) ratio in glial cells across time in culture (control n = 371; ln2PMMA n = 408). **(E)** Confocal images of YAP/TAZ immunostaining (green). **(F)** Linear regression analysis of YAP/TAZ Nuc/Cyto ratio in glial cells across time in culture (control, n = 274; ln2PMMA, n = 346). **(G)** Plot showing the inverse correlation between *Lmnb1* and *Lmna* gene expression, normalized to the mean *Lmna* expression level at E13, together with the expression of markers associated with RG to astrocyte lineage progression during mouse brain development. **(H)** Combined dot plot and heat map showing the percentage of cells expressing each marker (dot size) and the average expression level (color) across RG to astrocyte lineage cell populations in the mouse cerebral cortex at different embryonic (E) and postnatal (P) stages. Expression values were normalized to the maximum expression level of each gene (range 0-1). Data in (**G-H**) were obtained from Di Bella et al. (2021). Statistical significance of linear regressions was assessed by ANOVA (**p<0.01). Scale bars, (**A**) = 10 µm; (**C**-**D**) = 20 µm

To further contextualize the link between nuclear lamina remodeling and mechanosensitive signaling, we investigated whether mechanotransduction pathways are coordinately regulated during the RG to astrocyte lineage transition *in vivo*. To this end, we performed a meta-analysis of embryonic to early postnatal mouse cortical scRNA-seq datasets^43^ (**Fig. 2G-H**, **S1**). The RG to astrocyte transition was associated with a Lamin isoform shift, from high *Lmnb1* to high *Lmna* expression, accompanied by lineage-dependent changes in mechanosensitive ion channels (*Kcnk1/2/10, Trpc1, Pkd2, Tmem120a*), Ca²⁺ regulators (*Orai1, Tmco1, Tmem38a, Grina* and MCU components) and connexin-43 (*Gja1*). Although nuclear morphology cannot be directly assessed in scRNA-seq data, the coordinated regulation of lamina components and calcium-related genes is consistent with a shared mechanobiological program associated with RG lineage progression and the emergence of RG-associated cellular states.

### Mechanical cues reshape nuclear Ca²⁺ dynamics in RG-associated cell states

Because Ca²⁺ dynamics differ between RG and astrocytes *in vivo*^44–46^, we quantified Ca²⁺ activity in 3-4 DIV glial cultures loaded with Oregon Green™ BAPTA-1, AM (OGB). Measurements were obtained from regions of interest (ROIs) positioned withing the nuclei and correlated with Pax6 and Nestin expression (n = 529 nuclei; **Fig. 3A-B**), thereby primarily reflecting somatic/nuclear rather than whole-cell Ca²⁺ dynamics. Mean nuclear Ca²⁺ levels were significantly lower on ln2PMMA than on control substrates, irespectively of extracellular Ca²⁺ conditions (**Fig. 3C-D**). Stratification by marker expression revealed a consistent hierarchy of Ca²⁺ levels (Pax6+ > Nestin+/Pax6-> Nestin-/Pax6-). Notably, ln2PMMA selectively reduced Ca²⁺ levels in Pax6-negative nuclei, while largely preserving the elevated levels observed in Pax6+ nuclei (**Fig. 3E-G**). Spontaneous nuclear Ca²⁺ dynamics were also influenced by substrate topography. Silent nuclei were more frequent on ln2PMMA (42%) than on control (24%) substrates, and active nuclei on ln2PMMA displayed fewer and lower amplitude ΔF/F₀ events (**Fig. 3H, J-K**). Despite this global reduction in activity, Pax6+ nuclei maintained robust oscillatory behavior across substrates (**Fig. 3I**), indicating preservation of dynamic nuclear Ca²⁺ signaling within this population. Analysis of signal components further revealed substrate-dependent differences. Ca²⁺ activity on control substrates typically comprised both phasic and tonic components, whereas phasic events, previously associated with intracellular Ca²⁺ store release^45^, were markedly reduced on ln2PMMA across cell types. In contrast, tonic activity, linked to Ca²⁺ influx^45^, was comparatively preserved (**Fig. 3L-M**), becoming the predominant signaling mode in ln2PMMA cultures. Together, these results show that mechanical cues reorganize spontaneous nuclear Ca²⁺ signaling in a cell-state-dependent manner. Although ln2PMMA broadly reduces nuclear Ca²⁺ levels and phasic activity, Pax6+ nuclei retain distinctive oscillatory dynamics, suggesting that specific Ca²⁺ signaling states are associated with RG-like competence.

**Figure 3.**
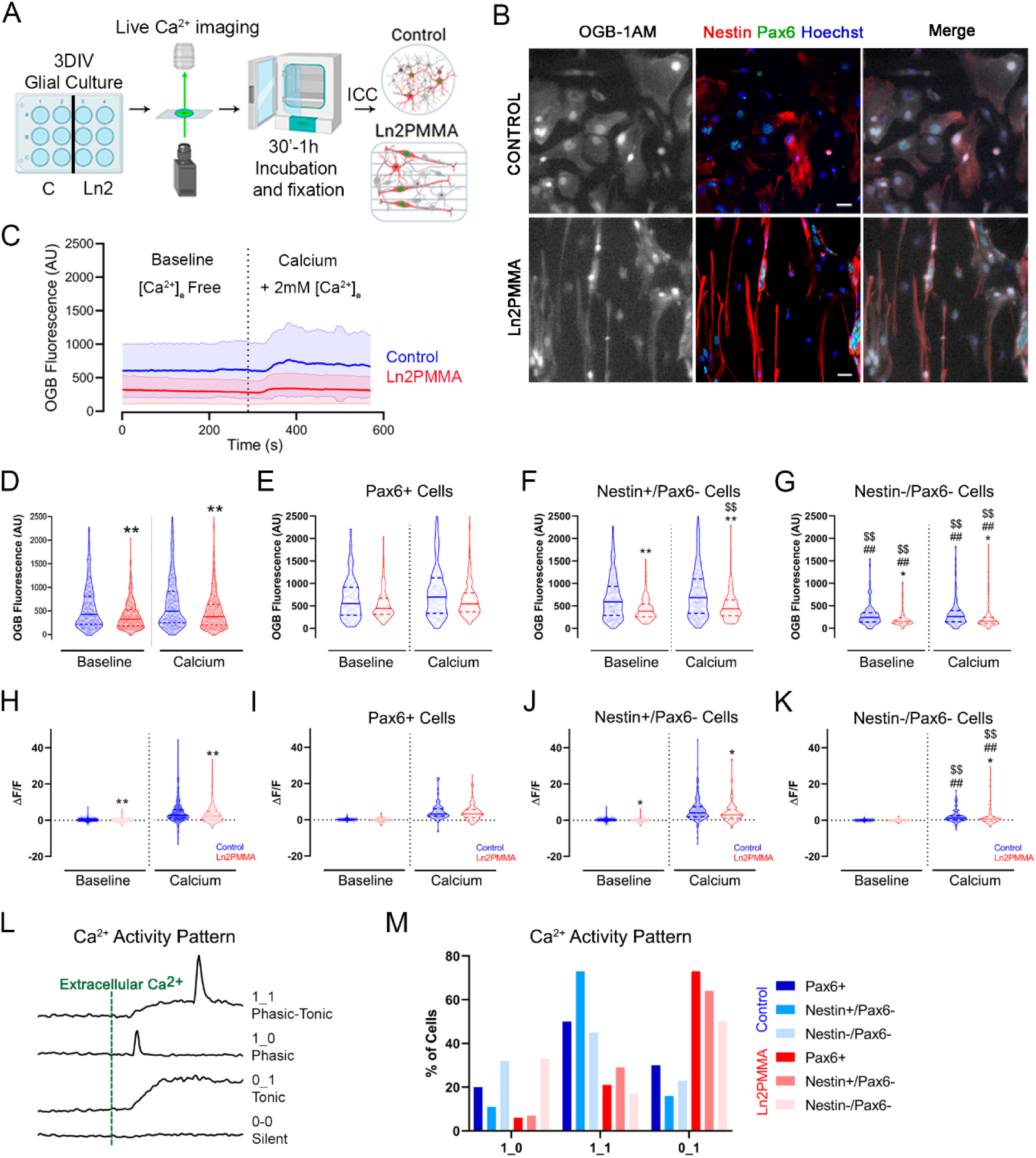
Ca^2+^ dynamics in glial cells grown on control and ln2PMMA substrates. **(A)** Experimental design for *in vitro* Ca^2+^ recordings. **(B)** Representative microscopy images of glial cells cultured for 3-4 DIV on control and ln2PMMA substrates, live-loaded with OGB (white), and immunostained after fixation for the RG-associated markers Pax6 (green) and Nestin (red). Nuclei were counterstained with Hoechst 33342 (blue). **(C)** Intracellular Ca²⁺ levels within nuclear ROIs were quantified using a background-subtraction method [F(t) – background] and are shown as mean signal ± 95% confidence interval (CI; shaded area) for control (blue, n = 257 cells) and ln2PMMA (red, n = 272 cells). Baseline corresponds to Ca²⁺-free extracellular solution, wereas “calcium” refers to extracellular solution containing 2 mM CaCl₂. **(D-G)** Violin plots showing average fluorescence intensity in **(D)** all nuclei corresponding to panel (**C**), (**E**) Pax6⁺ cells (control n = 63; ln2PMMA n = 91), **(F)** Nestin⁺/Pax6⁻ cells (control n = 125; ln2PMMA n = 106), and **(G)** Nestin⁻/Pax6⁻ cells (control n = 69; ln2PMMA n = 75). **(H-K)** Violin plots of spontaneous nuclear Ca^2+^ transients (mean ΔF/F_0_) under baseline and calcium conditions in **(H)** all glial cells, **(I)** Pax6+ cells, **(J)** Nestin+/Pax6-cells, and **(K)** Nestin-/Pax6-cells grown on control (blue) and ln2PMMA (red) substrates. **(L)** Schematic representation of the four patterns of Ca²⁺ activity identified above the ΔF/F₀ ≥ 3% threshold. **(M)** Distribution (%) of Ca²⁺ activity patterns among active Pax6⁺ cells, Nestin⁺/Pax6⁻, and Nestin⁻/Pax6⁻ glial cells grown on control (blue) and ln2PMMA (red) substrates. Scale bars = 25 µm. Four independent samples per condition. Statistical analyses were performed using the Kruskal-Wallis test with post hoc multiple comparisons. Differences between control and ln2PMMA within the same cell type: *p < 0.05, **p < 0.01; differences relative to Pax6⁺ cells within the same substrate: $$p < 0.01; differences relative to Nestin⁺/Pax6⁻ cells within the same substrate: ##p < 0.01.

### Calcium signaling contributes to RG-associated marker induction while nuclear remodeling persists independently of Ca²⁺ inhibition

To determine whether Ca²⁺ signaling relates to nuclear remodeling and/or substrate-induced gene expression changes, we transiently inhibited MSCs with GsMTx4 (1.22 µM) or CaMKII with KN93 (5 µM) for 3 h at 3 DIV (**Fig. 4A, D**). GsMTx4 is a non-selective inhibitor of Ca^2+^ influx through MSCs in response to membrane deformation^47^, thus predominantly affecting the mechanoelectrical tonic component of the Ca²⁺ signal. KN93 is a competitive inhibitor of CaMKII, a major mediator of Ca^2+^-dependent signaling^48,49^. Neither treatment affected cell viability (**Fig. S2**). Both inhibitors abolished substrate-dependent differences in marker expression, although through different mechanisms: GsMTx4 prevented Pax6 induction on ln2PMMA (**Fig. 4B**), and KN93 increased GFAP expression under the same conditions (**Fig. 4E**). These results indicate that Ca²⁺-dependent pathways are required for substrate-dependent Pax6 induction and other RG-associated marker responses. Despite these effects on lineage marker expression, neither inhibitor disrupted ln2PMMA substrate-induced nuclear elongation (**Table S1**), indicating that mechanical loading and nuclear remodeling persist under short Ca²⁺ inhibition. Both inhibitors reduced the Lamin A/C to B1 ratio on control substrates, shifting values toward those observed on ln2PMMA, although ln2PMMA consistently maintained the lowest ratios (**Fig. 4C, F**). These effects are consistent with known influences of Ca²⁺-dependent pathways on Lamin dynamics^50^ and suggest possible feedback between calcium signaling and nuclear architecture.

**Figure 4.**
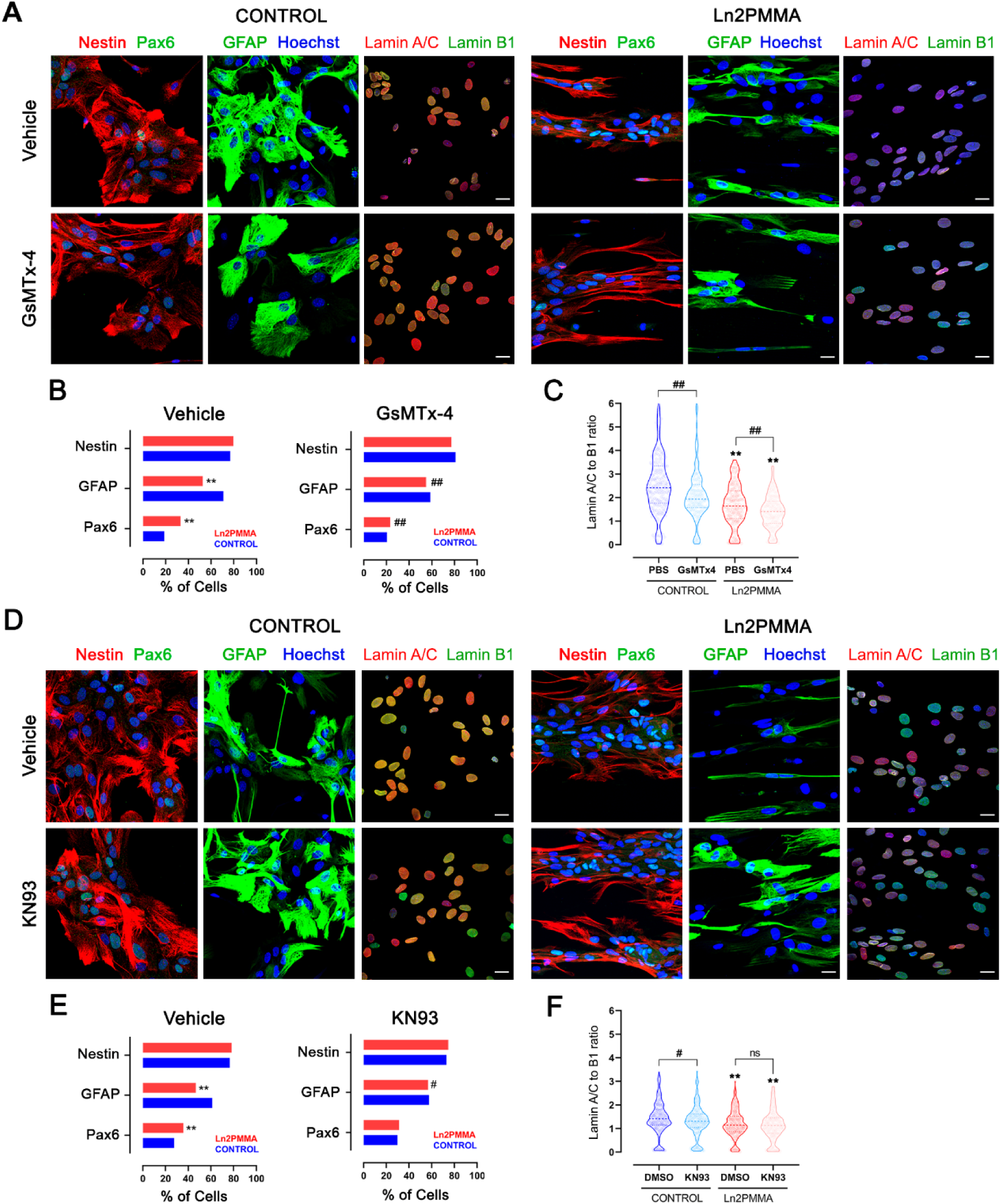
Pharmacological validation of MSCs involvement and Ca^2+^ signaling in the biomechanical induction of RG-like competence. **(A)** Confocal images of glial cells cultured on control and ln2PMMA substrates for 3 DIV and treated for 3 h with vehicle (PBS) or MSCs inhibitor GsMTx4 (1.22 µM). Cultures were immunostained for the RG-associated markers Nestin and Pax6, the astrocyte marker GFAP, and for Lamin A/C and Lamin B1. Nuclei were counterstained with Hoechst 33342 (blue). **(B)** Bar plots showing the percentage of cells expressing Nestin, Pax6 or GFAP in the cultures described in (**A**) (vehicle, n = 372; GsMTx4, n = 398). **(C)** Violin plots showing Lamin A/C to Lamin B1 ratios in the cultures described in (**A**). **(D)** Confocal images of glial cells cultured for 3 DIV on control and ln2PMMA substrates and treated for 3 h with vehicle (DMSO) or the CaMKII inhibitor KN93 (5 µM). Cultures were immunostained as in (**A**). **(E)** Bar plots showing the percentage of cells expressing Nestin, Pax6 or GFAP in the cultures described in (**D**) (vehicle, n = 483; KN93, n = 506). **(F)** Violin plots showing Lamin A/C to Lamin B1 ratios in the cultures described in (**D**). Scale bars = 20µm. Statistical analyses were performed using the Kruskal Wallis test followed by post hoc multiple comparisons. Differences between control and ln2PMMA under the same pharmacological condition: *p < 0.05, **p < 0.01. Differences between pharmacological treatments within the same substrate: #p < 0.05, ##p < 0.01. Data represent at least two independent biological samples per condition. Scale bar = 20 µm.

Short-term inhibition abolished substrate-dependent Pax6 induction while substrate-induced nuclear elongation persisted, indicating that early structural remodeling and RG-associated marker induction can be experimentally uncoupled.

Taken together, these findings indicate that mechanical and mechanoelectrical signaling can be differentially regulated. While Ca²⁺-dependent pathways are required for efficient Pax6 induction, substrate-induced nuclear remodeling persists under short-term Ca²⁺ inhibition. These observations support a model in which early mechanically induced structural remodeling and subsequent RG-associated marker induction represent separable processes. While these observations are consistent with a role for nuclear mechanics preceding RG-associated marker induction, we cannot exclude parallel, coordinated or bidirectional interactions between nuclear architecture and calcium signaling.

### Nuclear morphology captures features associated with RG-like competence

To determine whether nuclear morphology reflects RG-like competence, we analyzed whether simple geometric nuclear features could distinguish morphometric states associated with RG-like cell populations. Astrocyte cultures grown for 3 DIV on control or ln2PMMA substrates were stained for lineage markers, and nuclear area and eccentricity were extracted from 3,855 Hoechst-stained nuclei (**Fig. 5A-B**). Kernel density estimation revealed a significant shift toward higher nuclear eccentricity on ln2PMMA in Nestin+ (RG, immature astrocyte), GFAP+ (astrocyte), Pax6+ (RG), and Sox2+ (general stem cells) populations (p < 0.01), whereas NG2⁺ oligodendrocyte precursor cells and Gsx2+ intermediate progenitors were largely unaffected (**Fig. 5C; Table S2**). These findings indicate that mechanically induced changes in nuclear geometry occur selectively in cell populations associated with RG-like competence.

**Figure 5.**
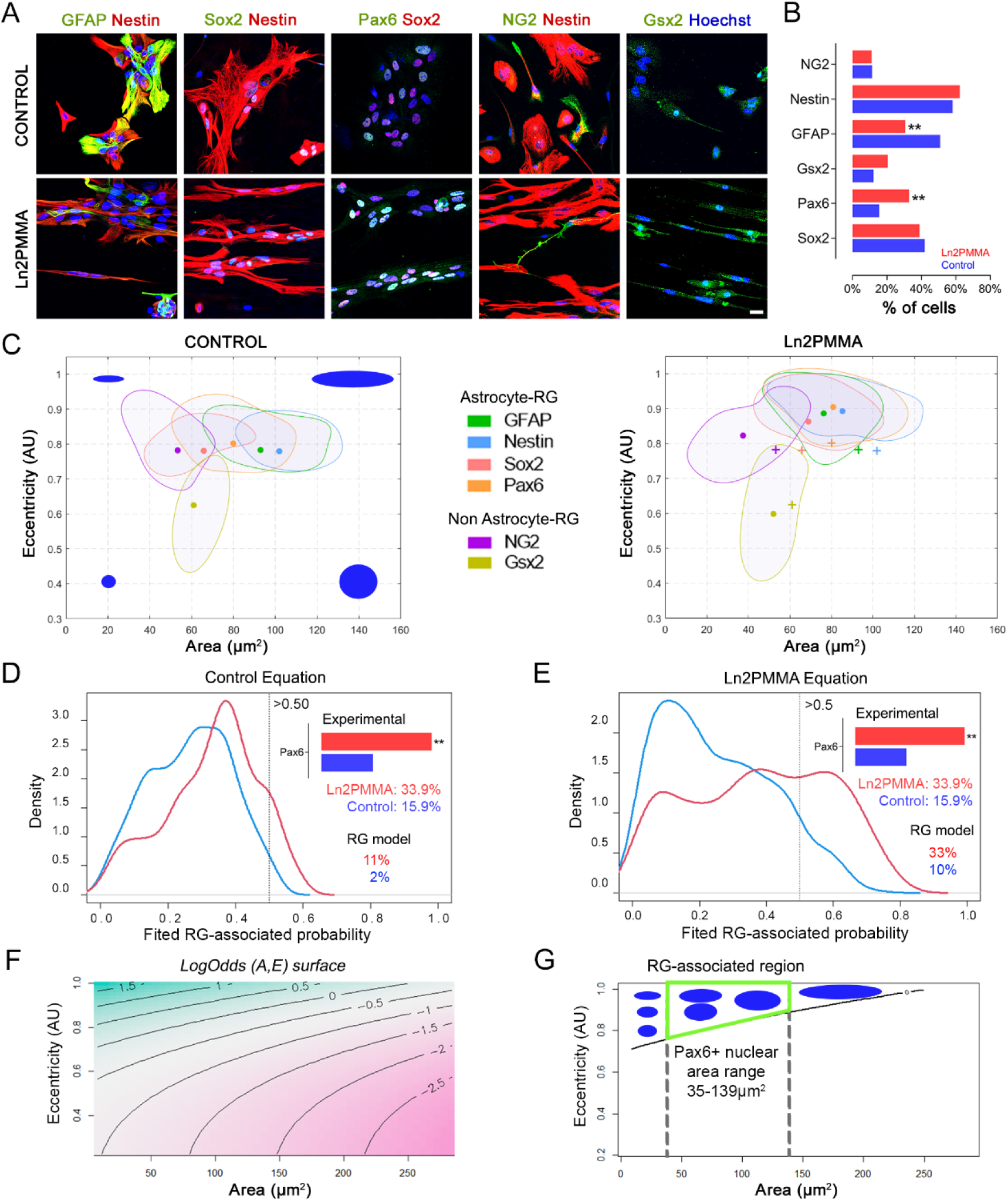
Generation of a statistical model capturing morphometric features associated with RG-like competence. **(A)** Representative images of primary cortical glial cells cultured for 3 DIV on control and ln2PMMA substrates and immunolabeled with cell type-specific markers for astrocytes (GFAP), immature astrocytes/RG (Nestin), RG-associated cells (Pax6), general stem cells (Sox2), and intermediate progenitors (NG2, Gsx2). Nuclei were counterstained with Hoechst (blue). **(B)** Bar plot showing the percentage of each cell type on control and ln2PMMA substrates. **(C)** Multivariate kernel density estimates showing the distribution of nuclear eccentricity (y-axis) and nuclear area (x-axis) for each cell type on control and ln2PMMA substrates. Centroids (dots) represent median values, and “+” symbols in the ln2PMMA plots indicate the corresponding centroids under control conditions. Blue shapes depict the reference area-eccentricity combinations. **(D-E)** Density plots of RG-associated probabilities predicted by the fitted models obtained by applying the **(D)** control-derived and **(E)** ln2PMMA-derived equations to the training dataset. Dashed lines indicate the probability threshold for RG-associated states (RGp = 0.5). Numbers indicate the percentage of cells with RGp above this threshold in each condition. Bar plots show the proportion of Pax6+ cells at 3 DIV in the experimental dataset. **(F)** Surface plot showing the LogOdds(A,E) surface as a function of nuclear area (x-axis) and eccentricity (y-axis). **(G)** Scatter plot showing F(0) level set of the LogOdds(A,E) surface (ascending black line), corresponding to the threshold for RG-associated states, together with representative nuclear morphologies (blue shapes). Vertical dashed lines indicate the nuclear area range of Pax6+ cells. The theoretical region corresponding to RG-associated morphometric states is highlighted in green. Chi-square test, **p<0.01. Scale bar = 10 µm.

To quantify these features, we built a logistic regression model in which RG-associated states were operationally defined based on marker expression (Pax6+ or Sox2+/Nestin+ cells). Thus, the model captures morphometric patterns associated with marker-defined RG-like populations rather than independently assigning cell identity. Because cell-cell interactions influence nuclear morphology and responses to substrate-derived mechanical cues^35,51^, we included cell density together with nuclear area and eccentricity. Generalized linear models with a logit link were fitted and iteratively simplified (Methods). The final model (AIC = 1300.33) achieved a modest but robust predictive performance (AUC overall 0.73; ln2PMMA 0.77; control 0.67), consistent with an interpretable low-dimensional description. Model coefficients (**Table S3**) indicated that substrate-dependent interactions, particularly those involving eccentricity, were the strongest contributors. Higher-order eccentricity terms (quadratic and cubic) were retained for both substrates, emphasizing the importance of nuclear elongation, whereas cell density contributed only through interactions with area and eccentricity, suggesting a modulatory role. Because the unified model retained substrate-dependent interaction terms, it could be expressed as two substrate-conditioned equations, corresponding to control and ln2PMMA conditions (**Table S3**, Methods). We next compared the predictions generated by each substrate-conditioned formulation with the observed marker-defined distributions. Using a predicted RG probability (RGp) threshold of 0.5, the control-conditioned equation underestimated the proportion of Pax6⁺ cells in both conditions (control: 2% estimated vs. 15.5% observed; ln2PMMA: 11% estimated vs. 32.9% observed; **Fig. 5D**). In contrast, the ln2PMMA-conditioned equation more closely matched the experimental values (control: 10%; ln2PMMA: 33%; **Fig. 5E**). We therefore used the ln2PMMA-conditioned formulation as the Radial Glia Mimetic (RGM) model. The RGM model represents a morphometric framework associated with RG-like competence under conditions that promote RG-associated marker induction.

Applying the RGM model to marker-defined subpopulations (**Fig. S3**) revealed high RG-associated probabilities for Pax6+ (alone or in combination with Nestin, GFAP or Sox2) and Nestin+/Sox2+ nuclei, with clear substrate dependence (ln2PMMA: 0.8-0.51; control: 0.22-0.15, depending on the marker combination) (**Fig. S3A, E**). In contrast, Nestin or GFAP expression alone did not reliably stratify these subpopulations, and combinations including NG2 or Gsx2 remained associated with low RG probabilities. These results indicate that marker-defined RG-associated states occupy distinct regions of morphometric space that depend on mechanical context.

To identify morphometric regions differentially associated with RG-like states across substrates, we compared the substrate-conditioned model equations using an odds-ratio transformation. The resulting LogOdds function depended only on nuclear area and eccentricity (**Table S3**), because density-dependent terms cancelled in the comparison. Plotting this function defined a two-dimensional representation of morphometric space (**Fig. 5F**), in which the LogOdds = 0 isoline (F(0)) marked the boundary where the predicted odds of RG association were equivalent between control and ln2PMMA conditions. Regions above this boundary corresponded to morphometric states for which the ln2PMMA-conditioned formulation predicted higher RG-associated odds than the control-conditioned formulation. Using the empirically observed range of Pax6⁺ nuclear areas (34.75-139 μm²) as a reference, we identified a region of morphometric space associated with RG-like competence (**Fig. 5G**), characterized by intermediate nuclear area and high eccentricity.

Evaluation on two independent experimental datasets derived from independent biological samples at 3 DIV (n = 608, corresponding to the 3 DIV condition shown in **Figure 1C**, and n = 1349, respectively) showed comparable distributions between RGM-predicted RG-associated frecuencies and observed Pax6+ frequencies (ln2PMMA: RGM RG-associated estimated 21-31%; Pax6+ 22.7-33%; control: RGM RG-associated estimated 6%; Pax6⁺ 11.2-18.8%). RGM-assigned probabilities were near zero for neuronal and cycling nuclei (**Fig. S4;** p ≤ 0.05) and remained close to control values (p ≤ 0.14) for glial cells grown on larger linear topographies (ln10PMMA), which do not substantially induce Pax6 expression^35^, consistent with the absence of RG-associated nuclear remodeling under these conditions. Together, these results indicate that nuclear geometry provides a low-dimensional, interpretable descriptor of morphometric states associated with RG-like competence. Importantly, the model assigns RG-associated probabilities based solely on morphometric features, without requiring direct marker measurements. As expected for a low-dimensional model, predictive performance is moderate but consistent across experimental conditions.

### Conserved nuclear morphometric state across contexts associated with RG-like competence

If the RGM model captures a competence state rather than a marker-defined RG identity, RGM-associated probabilities should emerge before peak Pax6 expression. We then assessed whether this RGM-defined nuclear signature is conserved across time and experimental contexts. We first analyzed temporal dynamics of nuclear morphometric states at the cohort level across defined time points (1-14 DIV) by comparing the proportion of nuclei with estimated RGp ≥ 0.5 to experimentally measured Pax6+ cells (**Fig. 6A).** Using the longitudinal dataset presented in **Figure 1B-C**, which spans the entire astrocyte-RG-astrocyte cycle^35^, we observed distinct temporal profiles for morphometric and molecular marker readouts. In control cultures, Pax6+ frequencies remained relatively stable (9.9-15.7%), while RGM estimates were consistently low (5-6%). This indicates that the model does not assign high RG-associated probabilities to the persistent Pax6+ population in control conditions. In ln2PMMA cultures, Pax6+ frequencies peaked at 3-7 DIV (22.7-21.6%), returning to control-like values at 1 and 14 DIV. In contrast, RGM-estimated RG-associated probabilities peaked earlier (1-3 DIV; 22-21%) and then declined (7-14 DIV; 15-12%). Because Pax6 expression was included during model training, this temporal offset suggests that the RGM model captures morphometric features that precede Pax6 induction. Importantly, RGM-associated probabilities peaked before maximal Pax6 expression, indicating that morphometric remodeling emerges earlier than RG-associated marker induction. This behavior is consistent with the existence of a competence-associated morphometric state preceding peak marker expression.

**Figure 6.**
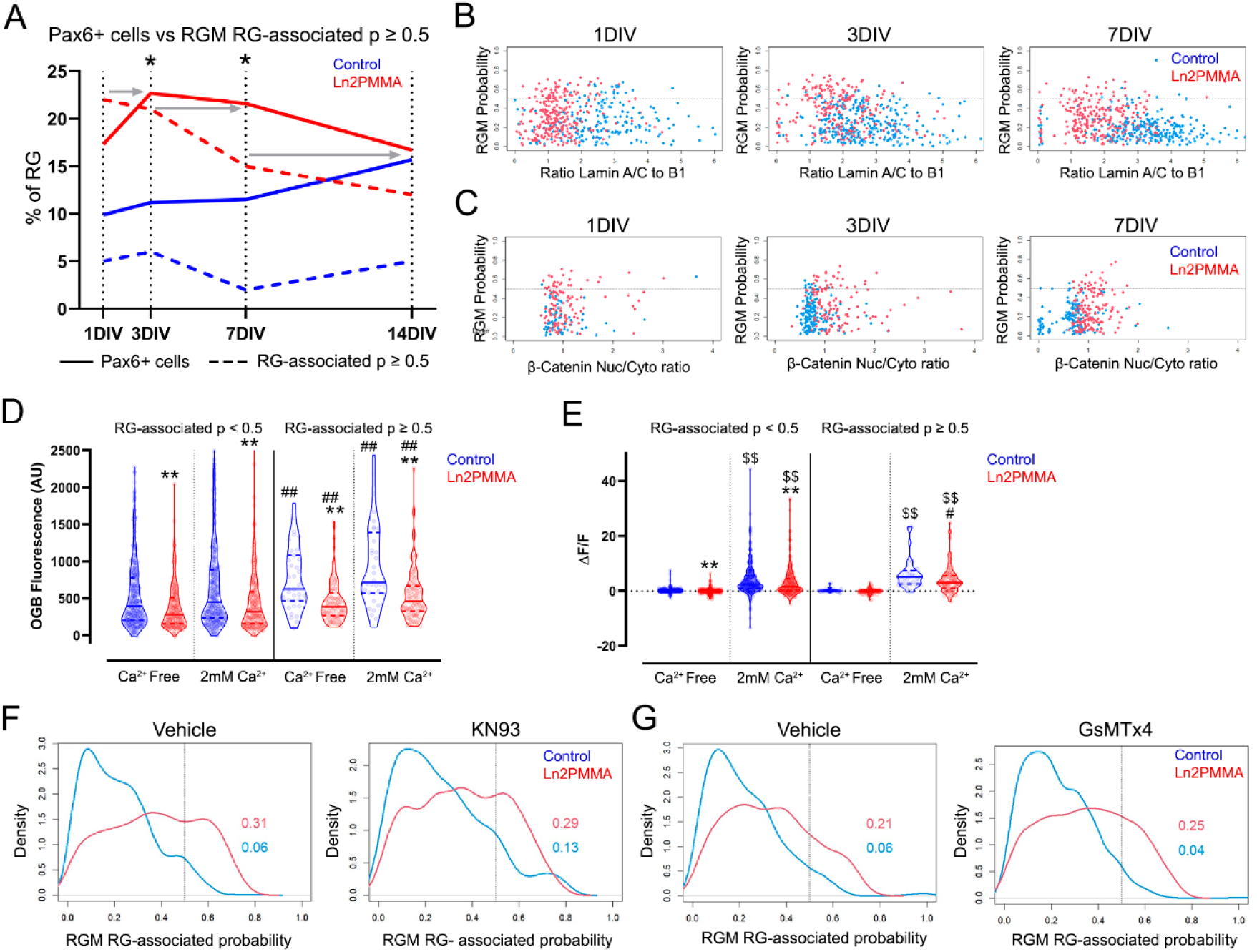
Conservation of a nuclear morphometric state associated with RG-like competence across contexts. **(A)** Comparison of experimentally quantified Pax6+ cell fractions (solid lines) and RG-associated fractions predicted by the RGM model (dashed lines) over time *in vitro*, using the dataset from Figure 1. Gray arrows indicate temporal offsets between experimental and model-predicted RG estimates. Statistical significance was assessed using a chi-square test (*p < 0.05). **(B)** Scatter plots showing RG-associated probabilities as a function of increasing Lamin A/C to B1 ratios from 1 to 7 DIV, using the dataset from Figure 2. The dashed line indicates the probability threshold for RG-associated states (RGp = 0.5). Cells cultured on control substrates are shown in blue, and cells cultured on ln2PMMA substrates are shown in red. **(C)** Scatter plots showing RGM-predicted RG-associated probabilities as a function of increasing nuclear to cytoplasmic (Nuc/Cyto) β-catenin ratios from 1 to 7 DIV, using the dataset from Figure 2. The dashed line indicates the RGp = 0.5 probability threshold. Control substrates are shown in blue and ln2PMMA sunstrates in red. **(D)** Violin plots showing basal nuclear Ca²⁺ levels under calcium-free and calcium-containing conditions in RGM-defined RG-associated (RGp ≥ 0.5) and non-RG-associated (RGp < 0.5) populations, using the dataset from Figure 3. **(E)** Violin plots showing spontaneous nuclear Ca²⁺ transients under calcium-free and calcium-containing conditions in RGM-defined RG-associated morphometric states (RGp ≥ 0.5) and lower-probability populations (RGp < 0.5), using the dataset from Figure 3. Statistical analysis was performed using a Kruskal-Wallis test with multiple comparisons and Benjamini-Yekultieli correction. **p < 0.01 indicates differences between control and ln2PMMA conditions within the same RGM RG-associated probability group; ##p<0.01 indicate differences between high RG-associated probability (RGp ≥ 0.5) and low RG-associated probability (RGp < 0.5) nuclei within the same substrate under either calcium-free or calcium-containing conditions; $$p < 0.01 indicates differences between absence and presence of extracellular Ca^+2^ within the same substrate and RG-associated probability group. Data were obtained from four independent samples per condition. **(F-G)** Density plots of RGM-predicted RG-associated probabilities showing probability distributions in cultures treated with KN93 (5 µM) **(F)**, GsMTx-4 (1.22 µM) **(G)**, and their corresponding vehicle controls, using the dataset from Figure 4. Dashed lines indicate the RGp = 0.5 probability threshold. Control substrates are shown in blue and ln2PMMA substrates in red. Numbers indicate the percentage of cells with RGp ≥ 0.5.

Having established that mechanically induced nuclear remodeling precedes and persists independently of Ca²⁺-dependent RG-associated marker induction, we next asked whether nuclear morphometric properties alone are associated with RG-like competence across contexts. To address this, we applied the RGM model to datasets quantifying Lamin A/C to B1 ratios, β-catenin localization, and nuclear Ca²⁺ activity. By analyzing the distribution of nuclei with RGM-estimated RGp ≥ 0.5 for different Lamin A/C to B1 ratios, we observed a trend toward lower ratios, although no specific Lamin A/C to B1 ratio was significantly associated with high RG probability (**Fig. 6B**). Likewise, nuclear/cytoplasmic β-catenin ratios did not independently stratify RG-associated states (**Fig. 6C**), indicating that lamina remodeling and β-catenin nuclear retention are associated with, but do not define, this morphometric state.

In contrast, applying the RGM model to nuclear Ca²⁺ datasets revealed that nuclei with higher RG-associated probabilities (RGp ≥ 0.5) consistently displayed elevated basal nuclear Ca²⁺ levels and more frequent nuclear Ca²⁺ transients than nuclei with low RG-associated probabilities (RGp < 0.5), independent of substrate or extracellular Ca²⁺ availability (**Fig. 6D-E**). This pattern mirrors spontaneous somatic Ca²⁺ behavior of RG in the embryonic cortical ventricular zone^44^ and differs from the low somatic spontaneous activity profiles typical of astrocytes^45^. Application of the RGM model to pharmacological datasets further showed that short-term inhibition of mechanosensitive ion channels or CaMKII did not reduce the fraction of cells occupying the RG-associated morphometric state (0.21 to 0.25 under GsMTx4; 0.31 to 0.29 under KN93), despite the loss of Pax6 induction (**Fig. 6F-G**). Pharmacological uncoupling of RGp and Pax6 expression further supports a temporal sequence in which mechanically associated morphometric states emerge before efficient RG-associated marker induction.

Together, these findings indicate that morphometric properties delineate a state associated with RG-like competence within a mechanical context. In this framework, Ca²⁺ signaling is required for efficient RG-associated marker induction, as reflected by Pax6 expression, but is not necessary for the establishment or maintenance of the underlying morphometric state.

### RG-associated nuclear morphometric state is conserved across species

We therefore asked whether this mechanically encoded nuclear state is preserved in native developmental contexts and across species. To examine whether nuclear morphometric states identified *in vitro* are also observed *in vivo*, we applied the substrate-independent LogOdds function, based solely on nuclear area and eccentricity, to embryonic day 18 (E18) mouse cortical sections immunostained for Pax6, using the same empirically defined nuclear area range established *in vitro* (n = 891 nuclei). As expected, the proportion of nuclei falling within the RG-associated region was highest in the ventricular zone (VZ), intermediate in the subventricular zone (SVZ), and lowest in the intermediate zone (IZ) (**Fig. 7A**). These distributions closely mirror established regional compositions and indicate that nuclear morphometric states observed *in vitro* are spatially organized *in vivo*, rather than arising from specific culture conditions.

**Figure 7.**
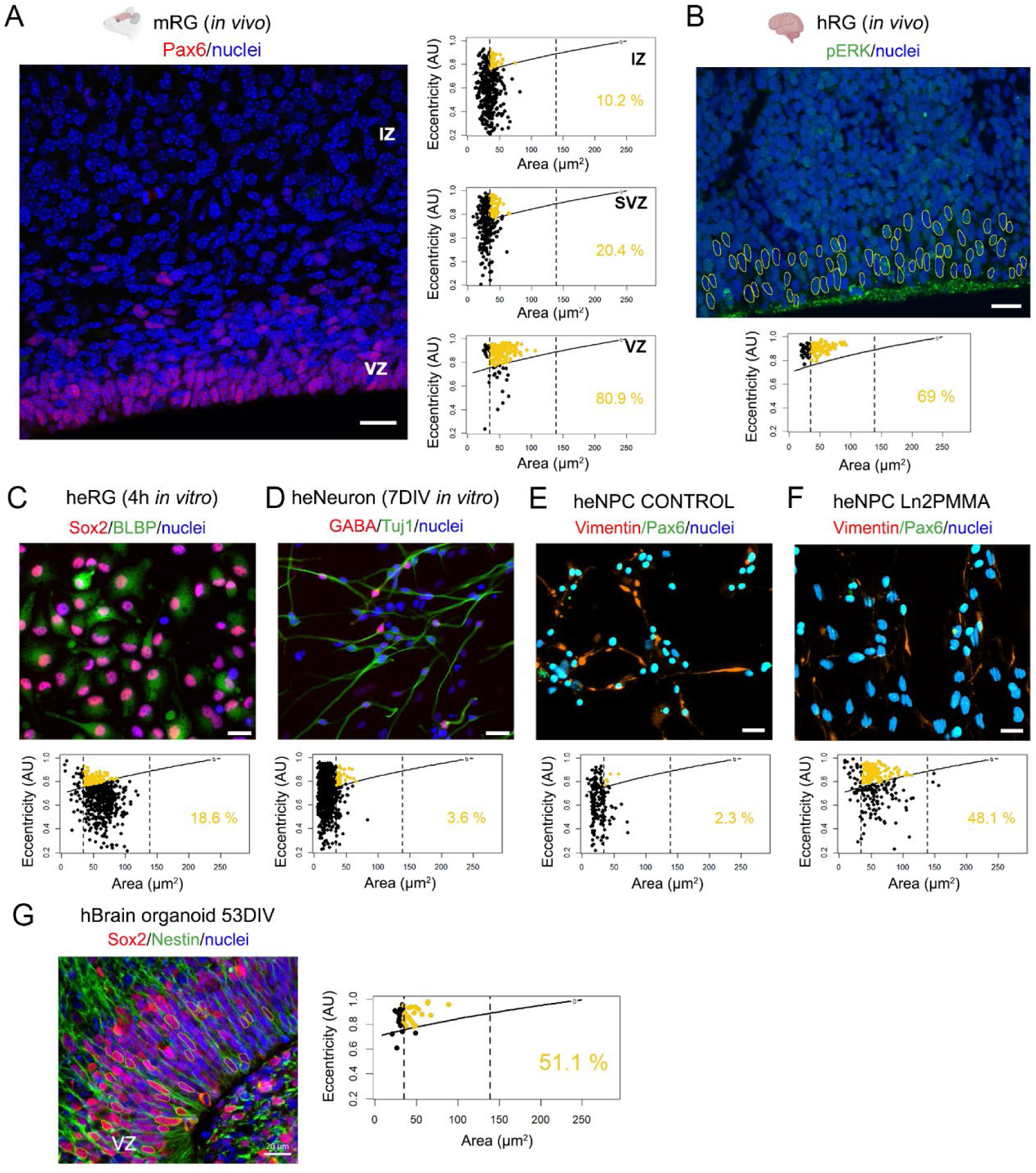
Conservation of a nuclear morphometric state associated with RG-like competence in mouse and human cortex. **(A)** Confocal image of a coronal section of the mouse cerebral cortex at embryonic day 18 (E18) immunostained for Pax6 (red) with nuclei counterstained with Hoechst 33342 (blue). Corresponding scatter plots on the right depict the LogOdds (A,E) function at F(0) (black curve) for distinct cortical regions. Yellow dots and percentages indicate nuclei (VZ, n = 194; SVZ, n = 313; IZ, n = 384) falling within the RG-associated morphometric region defined by the RGM model. Vertical dashed lines denote the nuclear area range observed in Pax6+ cells from mouse glial cultures (A_min_ = 34.75 µm², A_max_ = 139 µm²). Statistical analysis: Kruskal-Wallis test followed by post-hoc multiple comparisons analyses. Differences between control and ln2PMMA conditions: *p<0.05, **p<0.01; differences between cell types or treatments: #p<0.05, ##p<0.01. VZ, ventricular zone; SVZ, subventricular zone; IZ, intermediate zone. **(B)** Representative fluorescence micrograph of a coronal section of human fetal cerebral cortex at gestational week 10 (GW10), immunostained for phosphorylated ERK (pERK; green, with nuclear signal highlighted) and counterstained with DAPI (blue). The corresponding scatter plot is shown below (n= 90 nuclei). **(C)** Human radial glia (hRG) cultures derived from the tissue shown in (**B**) and cultured for 4 h on a flat control substrate, immunostained for Sox2 (red) and BLBP (green), with nuclei counterstained with DAPI (blue). The corresponding scatter plot is shown below (n= 585 nuclei). **(D)** Human RG-derived neurons after 7 days *in vitro* (7 DIV), immunostained for GABA (red) and Tuj1 (green), with nuclei counterstained with DAPI (blue). The corresponding scatter plot is shown below (n= 1014 nuclei). **(E-F)** Human induced neural progenitor cells (hiNPCs) cultured on a control substrate (Matrigel-coated; n = 214 nuclei) **(E)** or on ln2PMMA substrates (n = 266 nuclei) **(F)**, immunostained for vimentin (red) and Pax6 (green). Nuclei were counterstained with Hoechst 33342 (blue). The corresponding scatter plots are shown below. **(G)** Representative confocal image of a section from a 53 DIV human brain organoid, immunostained for Sox2 (red) and Nestin (green), with nuclei counterstained with Hoechst 33342 (blue). The corresponding dispersion plot is shown on the right (n = 48 nuclei). Scale bars = 20µm.

To assess whether this nuclear morphometric signature is conserved across species, we analyzed histological sections from fetal human cortex, as well as *in vitro* RG cultures derived from human tissue at 10-15 gestational weeks (material from refs.^52,53^). In fetal cortical sections, putative human RG (hRG) were identified by their characteristic location within the VZ and by pERK expression, consistent with the requirement of ERK signaling for RG maintenance^54^ (n = 90 nuclei). Notably, 69% of hRG nuclei occupied the RG-associated region of the morphometric space defined by the model (**Fig. 7B**), closely matching the distribution observed in mouse tissue. These findings indicate that the morphometric features associated with RG-like competence are across species and developmental contexts and can be captured using the RGM framework.

We next asked whether mechanical disruption of apical and basal anchorage during tissue dissociation alters nuclear geometry. Primary hRG, purified by immunomagnetic isolation, were imaged at 4 hours and 7 DIV to monitor their progression toward neuronal differentiation^52,53,55^. After only 4 hours on standard flat poly-D-lysine (PDL) substrates, hRG rapidly shifted in the morphometric space, with the proportion of nuclei within the RG-associated region decreasing from 69% *in situ* to 18.6% (n = 585 nuclei) (**Fig. 7C**), consistent with rapid mechanical relaxation following detachment. By 7 DIV, only 3.6% of nuclei (n = 1014 nuclei) remained within the RG-associated region, indicating that hRG-derived neurons occupy a nuclear morphometric state distinct from that of *in situ* RG (**Fig. 7D)**. These observations highlight the sensitivity of nuclear morphometric states to microenvironmental mechanical conditions and their dynamic reorganization during differentiation.

We then evaluated whether substrate anisotropy, an emergent biophysical property of the native RG niche, could bias nuclear states in early human neural progenitors. Using cultures of human stem cell-derived neural progenitor cells (hNPC), we found that Pax6⁺ nuclei on flat substrates exhibited broad dispersion across the morphometric space, with only 2.3% (n = 214 nuclei) falling within the RG-associated region (**Fig. 7E**). In contrast, nearly 50% of hNPC nuclei (n = 266 nuclei) cultured on ln2PMMA (**Fig. 7F**), and 51.1% of nuclei (n = 48) in the ventricular zone of brain organoids generated from iPSCs (**Fig. 7G**) occupied this region. These results indicate that microscale anisotropy of the niche biases nuclear states toward a morphometric region associated with RG-like competence.

Together, these observations indicate that RG-associated morphometric states are preferentially observed under conditions of anisotropic mechanical organization, whether imposed by tissue architecture, substrate topography, or apical-basal polarity. This supports a conserved mechanobiological framework, modulated by niche mechanics, in which specific regions of nuclear morphometric space are consistently associated with RG-like competence across developmental systems and species.

Together, the temporal emergence of nuclear remodeling, the earlier peak of RGM-assigned probabilities relative to Pax6 expression, and the preservation of RGM-associated states after KN93 or GsMTx4 treatment support a model in which mechanical cues establish an early morphometric state associated with RG-like competence, whereas calcium-dependent pathways are required for efficient RG-associated marker induction.

## DISCUSSION

Our findings support the existence of a morphometric state associated with RG-like competence that emerges before peak RG-marker expression and persists independently of calcium-dependent RG-associated marker induction. The results presented here support a working model in which anisotropic mechanical cues first establish a morphometric state associated with RG-like competence, whereas calcium-dependent pathways are required for efficient induction of RG-associated markers.

RG cells play a central role in cortical development, generating neurons and glial cells while providing a scaffold for neuronal migration^1,2^. Despite this central role, the mechanobiological mechanisms underlaying RG-associated states remain incompletely understood^6–8^. Here, we used the term RG-like competence to describe the acquisition of RG-associated molecular markers, including Pax6, Sox2, and Nestin, together with morphometric features characteristic of radial glia, without directly assessing functional behavior. In this context, we define a mechanical state as a coordinated configuration of cellular and nuclear morphology, local cellular context, and associated signaling activity. Our results identify a conserved nuclear architecture associated with RG-like competence across diverse biological contexts. Rather than representing a passive correlate of cell state, this nuclear architecture may reflect an early mechanobiological component associated with the acquisition of RG-like competence.

Mechanical cues imposed by anisotropic substrates reshape the soma and nucleus within hours of adhesion, well before detectable changes in lineage marker expression. The early reduction in Lamin A/C to B1 ratios, sustained nuclear elongation, and stable nuclear alignment induced by ln2PMMA are consistent with a mechanically softened nucleus, dynamic chromatin organization^56^ and a directionally constrained lamina state. These features emerge within the time window reported for mechanosensitive NSC lineage commitment (2–3 days after adhesion^57^), supporting the view that nuclear architecture acts as a mechanosensitive integrator of topographical cues. More broadly, our results suggest that mechanical regulation emerges from the combined effects of substrate anisotropy and cell-cell interactions rather than from any single physical parameter.

Pax6 is a key transcription factor associated with RG identity and a direct downstream target of the Wnt/β-catenin pathway^41^. The sustained nuclear accumulation of β-catenin observed under anisotropic conditions provides a plausible link between nuclear mechanics and transcriptional regulation, consistent with reported interactions between Pax6 and β-catenin in RG self-renewal and differentiation^41,58,59^. These observations support a model in which mechanically induced nuclear configurations stablish a permissive morphometric state that precedes and facilitates RG-associated marker induction. Notably, the absence of detectable changes in YAP/TAZ localization suggests that topography-driven responses may engage signaling pathways distinct from canonical stiffness-dependent mechanotransduction. In parallel, lamina remodeling may contribute to RG-associated marker induction by modifying chromatin organization and accessibility, potentially including the Pax6 locus through interactions with chromatin-remodeling complexes such as BAF^60^.

Although nuclear architecture is associated with a mechanically induced RG-competent state, substrate topography also generates asymmetric forces that activate mechanosensitive ion channels. The relationship between nuclear architecture and calcium signaling is likely coordinated and potentially bidirectional, rather than strictly hierarchical. Our inhibition experiments show that Ca²⁺-dependent pathways are required for Pax6 induction, as short-term blockade of mechanosensitive ion channels or CaMKII completely abolished substrate-dependent Pax6 expression while preserving nuclear elongation. These findings indicate that structural remodeling and calcium-dependent transcription can be partially uncoupled while remaining functionally coordinated. Within this framework, mechanical cues bias cells toward an RG-like competent state, whereas calcium signaling is required for efficient induction of RG-associated markers. The association between RG-like nuclear geometry and elevated nuclear Ca²⁺ dynamics, including increased transient frequency, further supports a model in which calcium activity contributes to RG-associated marker induction without fully determining the underlying mechanically associated state. Together, these observations support a model in which cells occupy a continuous morphometric space shaped by mechanical cues, where states associated with RG-like competence can be partially uncoupled from Ca²⁺-dependent RG-associated marker induction.

A long-standing challenge in neural lineage classification is that canonical markers such as Sox2, Pax6, Nestin, and GFAP are broadly expressed across progenitor and glial populations^61–64^. Although machine learning approaches can classify cell states from morphology^65–69^, they often provide limited mechanistic interpretability. Our interpretable RGM model addresses this limitation by identifying low-dimensional nuclear morphometric states, defined by area, eccentricity, and local density, that are associated with RG-like competence. Notably, these morphometric states are conserved across mouse and human ventricular zone RG and are rapidly lost upon dissociation-induced relaxation of tissue architecture. Conversely, microscale anisotropy reinstates RG-associated nuclear morphology in human neural progenitors, indicating that nuclear architecture responds dynamically to the mechanical organization of the microenvironment. Together these findings support the view that RG-like cells, like many stem and progenitor populations, are intrinsically mechanosensitive^26,57,70–72^, and identify nuclear architecture as a dynamic and interpretable readout of microenvironmental mechanical cues. Rather than imposing an artificial cellular state, microtopography appears to stabilize a nuclear configuration that is mechanically associated with RG identity and is also observed *in vivo*.

An important question arising from these findings is whether nuclear morphometric states actively contribute to the acquisition of RG-like competence or instead represent a quantitative manifestation of the underlying mechanobiological processes. Our data support the existence of a mechanically associated morphometric state that precedes peak RG-associated marker induction, but establishing its causal role will require experimental approaches capable of modulating nuclear architecture independently of substrate-derived cues. Combining targeted perturbations of nuclear organization with quantitative mapping in morphometric space may help determine whether specific shifts in nuclear configuration are sufficient to influence RG-associated transcriptional programs. Addressing this question will be critical for determining whether nuclear morphometric states are simply readouts of mechanical organization or constitute an active intermediate layer linking tissue mechanics to RG-associated gene regulation.

Several limitations should be considered. First, nuclear architecture was not independently manipulated, and therefore its causal contribution to RG competence cannot be formally established. Second, our analyses rely primarily on 2D nuclear descriptors, whereas RG nuclei *in vivo* are embedded within complex three-dimensional tissue architectures. Third, the morphometric model was trained using marker-defined states and exhibits moderate predictive performance, capturing a state associated with RG-like competence rather than providing an independent definition of cell identity.

Together, our findings support a model in which mechanically induced morphometric states are associated with RG-like competence across developmental systems. In this context, microtopography may act by stabilizing nuclear configurations that are also observed *in vivo*. More broadly, these results identify nuclear architecture as an interpretable intermediate layer linking microenvironmental organization to RG-associated transcriptional programs. This framework provides a basis for understanding how physical and biochemical cues are integrated during neural development.

## METHODS

### Ethical statement

All animal procedures were approved by the Ethical Committee for Animal Experimentation of the University of Barcelona and the Generalitat de Catalunya (protocol n°: 9656) and were conducted in accordance with Spanish and European Union regulations.

Human induced pluripotent stem cell (iPSC) line 18a and embryonic stem cell (ESC) line HUES64 (Harvard Stem Cell Institute) were transferred to University of Barcelona-IDIBELL following approval by the Generalitat de Catalunya Health Department and under a Material Transfer Agreement (MTA n° A5700; execution form C21_177) (principal investigator J.A. Ortega).

Human fetal brain tissue samples were obtained and processed in Zecevic’s laboratory (University of Connecticut Health Center) by J.A. Ortega. Samples were originally sourced from Advanced Bioscience Resources (ABR) and Human Developmental Biology Resource (HDBR), following written informed parental consent and approval by the corresponding ethics committees^52,53^. All procedures involving human tissue were conducted in accordance with institutional guidelines, the Declaration of Helsinki, and University of Connecticut regulations.

### Mouse cortical glial primary cultures

Cortical glial cultures were prepared from newborn (P0-P1) Swiss albino mice of both sexes (>100 animals from at least 10 independent litters). Following brain extraction, cerebral cortices were dissected free of meninges in dissection buffer consisting of PBS supplemented with 0.6% of glucose (Sigma) and 0.3% bovine serum albumin (BSA; Sigma). Tissue was enzymatically digested in 5 ml trypsin (Life Technologies) supplemented with 500μl of DNAse I (Sigma) for 10 min at 37°C. Following digestion, cortices were mechanically dissociated in growth medium (GM), consisting of Dulbecco’s Modified Eagle Medium (DMEM, Biological Industries) supplemented with 10% normal horse serum (NHS, GIBCO), 1% penicillin-streptomycin (P/S; Biological Industries) and 1% L-glutamine (Biological Industries). After centrifugation and resuspension, passage 0 (Ps0) cells were plated into culture flasks and maintained for 14 days. Astrocyte-enriched cultures (hereafter referred to as astrocyte cultures) were obtained by shaken mixed glial cultures overnight at 37°C, followed by a medium change one day before the start of the experiment. This procedure reduces contaminating microglia and oligodendrocyte progenitor cells and was adapted from^73^. For all experiments, Ps0 astrocyte cultures were dissociated with trypsin, counted, and replated onto the indicated substrates (borosilicate glass coverslips or ln2PMMA) at a density of 2-4 x 10^4^ cells/cm^2^ in 24-well plates, depending on the experiment. Cells were maintained for 1-14 days in vitro (DIV) in experimental medium (EM), consisting of Neurobasal™ (Gibco) supplemented with 3% NHS, 1% P/S, and 1% L-glutamine. Because ln2PMMA substrates are hydrophobic and were used without additional surface coating, the cell suspension (60 µl) was carefully distributed across the substrate surface and allowed to adhere for 20-30 min at 37°C before addition of the culture medium (final volume: 500 µl per well).

### Human pluripotent stem cells, neural progenitors, and brain organoids

Human induced pluripotent stem cell (iPSCs) line 18a and embryonic stem cell (ESC) line HUES64 (Harvard Stem Cell Institute) were maintained in mTeSR™ medium (STEMCELL Technologies, #85850) on Matrigel-coated plates (Thermo Fisher scientific, BD354277) and fed daily. For thawing and passaging, cells were resuspended in DMEM/F-12 (Gibco, #11574546) and cultured in mTeSR supplemented with 10 µM ROCK inhibitor (Tocris, #1254) for 24-48 h. Routine passaging was performed using 0.5 mM EDTA, whereas Accutase (Innovative Cell Technologies, AT-104) was used for dissociation of differentiated populations.

Human neural progenitor cells (hNPCs) were derived from iPSC or ESCs as previously described^74^. Briefly, pluripotent cells were dissociated using Accutase and replated onto Matrigel-coated plates in mTeSR medium supplemented with ROCK inhibitor. Cells were allowed to reach approximately 70% confluency (day 0), after which the medium was replaced with N2/B27 differentiation medium consisting of DMEM/F-12 supplemented with 1x N2 (Gibco, #12013479), 1xx0023;11530536), and 20 ng/ml recombinant human basic fibroblast growth factor (bFGF; Fisher Scientific, PHG0264). Neural induction was promoted by addition of dual-SMAD inhibitors (0.1 µM LDN193189 and 10 µM SB431542) for 6 days, with daily medium changes. From day 7 onward, laminin was added at a final concentration of 1 µg/ml. On day 12, neural rosettes were isolated using STEMdiff™ Neural Rosette Selection Reagent (STEMCELL Technologies, #05832) according to the manufacturer’s instructions. Cells were subsequently dissociated and replated onto Matrigel-coated plates or ln2PMMA substrates and maintained in NPC medium consisting of DMEM/F-12 supplemented with N2 and B27 for subsequent morphometric and immunofluorescence analyses.

Brain organoids were generated from 18a iPSCs following a previously described protocol^75^. Cells were dissociated using Accutase and seeded into a 96-well plate (9000 cells/well) in mTeSR supplemented with ROCK inhibitor (1:1000) to form embryoid bodies (EBs). After 48h, ROCK inhibitor was removed once EBs reached >350 μm in diameter and displayed a compact morphology. Cultures were maintained in mTeSR until EBs reached 500-600 μm (∼5 days). EBs were then transferred to neural induction medium consisting of DMEM/F-12, supplemented with 1% GlutaMAX™, 1% N2, 1% penicillin/streptomycin, 1% non-essential amino acids (NEAA), 1 μg/mL heparin (H3149-10KU, MERCK), with half-medium changes every 2 days. Following neuroepithelium formation (days 9-10), organoids were transferred to 60 mm Petri dishes (10-15 organoids per dish) in differentiation media without vitamin A (day 0) and maintained under static culture conditions. After 4 days, vitamin A was added and organoids were transferred to an orbital shaker. Differentiation medium consisted of DMEM/F-12 and Neurobasal™ (1:2) supplemented with 1% GlutaMAX™, 1% B27 with vitamin A, 1% N2, 1% penicillin/streptomycin, 1% NEAA, 10μg/mL insulin, and 2-mercaptoethanol (1:100). Cultures were maintained at 37°C throughout the differentiation period.

### Human fetal tissue samples and human RG cultures

Fetal brains included in this study (from two fetuses at 10 and 15 gestational weeks respectively) showed no signs of neurological disease upon detailed neuropathological examination. Developmental stage was determined from gestational age and ultrasound measurements. Tissue was collected in oxygenated Hank’s Balanced Salt Solution (HBSS; Invitrogen) and transported on ice. Following dissection, cortical tissue was mechanically and enzymatically dissociated using 0.025% trypsin-EDTA (Invitrogen) supplemented with DNase I (2 mg/mL, Sigma-Aldrich). Dissociated cells were cultured in poly-D-lysine-coated flasks (Sigma-Aldrich) containing roliferation medium (PM), consisting of DMEM/F12 (Invitrogen), supplemented with B27 (Invitrogen), basic fibroblast growth factor (bFGF; 10 ng/mL; Peprotech), epidermal growth factor (EGF; 10 ng/mL; Millipore), and penicillin/streptomycin (Invitrogen). After expansion, human radial glial cells (hRG) were immunomagnetically isolated using MACS® technology with anti-CD15-coated microbeads (Miltenyi Biotec), as previously described^52,53^. Purified hRG cells were plated onto poly-D-lysine-coated substrates and maintained in PM for 1-3 days prior to analysis or differentiation assays. For differentiation assays, bFGF and EGF were omitted from the culture medium.

Immunocytochemistry of cultured hRG cells and immunohistochemistry of fetal tissue sections were performed as described in the Immunofluorescence section below.

### PMMA micropatterning, characterization, and sterilization

Polymethyl methacrylate (PMMA) sheets (125 µm thickness) were obtained from Goodfellow Ltd. (UK). Ln2PMMA substrates were fabricated by the Nanolithography Laboratory of the Institute of Microelectronics of Barcelona (IMB, CNM-CSIC). Ln2PMMA micropatterns consisted of linear grooves 2 µm in width and 1 µm in depth, generated by nanoimprint lithography (NIL) and characterized as previously described^34,76^. This geometry was selected because it reproduces key anisotropic features of the RG niche and has previously been shown to promote RG-like transitions in astrocyte cultures^34,35^.

For cell culture experiments, ln2PMMA substrates were cut to fit in 24-well culture plates, sonicated for 10 min in sterile water, sterilized in 70% ethanol for 15-20 min, thoroughly rinsed in sterile water for 20 min, air-dried, and used without additional surface coating. Borosilicate glass coverslips were used as control substrates, as no significant differences in cell morphology or RG-associated marker expression were observed among non-patterned PMMA, glass, and standard tissue culture plastic, as previously reported^34^.

### Pharmacological treatments

To perturb mechanosensitive and calcium-dependent signaling pathways, glial cultures at 3 days in vitro (DIV) were treated for 3 h with the Ca^2+^/calmodulin-dependent protein kinase II (CaMKII) inhibitor KN93 (5 µM), or the non-selective mechanosensitive cation channel inhibitor GsMTx-4 (1.22 µM) on both control and ln2PMMA substrates. The 3 h treatment period was selected to assess acute effects of pathway inhibition while minimizing secondary changes associated with long-term culture responses. Corresponding vehicle controls were included for each treatment condition (DMSO for KN93 and PBS for GsMTx4). No serum deprivation was performed prior to treatment. Following treatment, samples were fixed and processed for immunofluorescence as described below. Data were obtained from two independent biological experiments. The total number of cells analyzed per condition was: DMSO (n = 1,832), KN93 (n = 1,931), PBS (n = 1,349), and GsMTx4 (n = 1,727). The reported cell numbers correspond to pooled observations across substrate conditions.

### Calcium Imaging

Glial cultures at 3-4 DIV were mounted in a microfluidic chamber (Open Diamond Bath Imaging Chamber for Round Coverslips; Warner Instruments) and loaded with the calcium-sensitive dye Oregon Green BAPTA-1 AM (OGB-1 AM, Life Technologies; 10 μM) in extracellular medium for 30 minutes at 37°C and 5% CO_2_. The bath solution consisted of 140 mM NaCl, 5.4 mM KCl, 1 mM MgCl₂, 10 mM HEPES, and 10 mM glucose, adjusted to pH 7.4. Recordings were performed under calcium-free conditions followed by supplementation with 2 mM CaCl_2_. Intracellular Ca^2+^ activity was monitored for 10 minutes, including a 2-min calcium-free baseline period followed by recording in the presence of extracellular calcium.

Imaging was performed using an Olympus IX71 inverted microscope equipped with an XLUMPLFLN 20XW water-immersion objective (20x/1.0; Olympus). OGB-1 AM was excited at 488 nm (50 ms exposure) using a Polychrome V light source (Till Photonics) with a Xenon Short-Arc lamp (Ushio), and a 505 nm dichroic mirror (Chroma Technology). Fluorescence emission was collected through a D535/40 nm emission filter (Chroma Technology) and detected using a C9100-13 EM-CCD camera (Hamamatsu). Images were acquired at 1 Hz at room temperature.

Following calcium imaging, the imaged area was marked on the coverslip. Samples were subsequently washed and maintained in extracellular medium at 37°C for 30-60 min prior to fixation and preparation for immunocytochemistry (see Immunofluorescence section below). For each substrate condition, eight samples were initially processed (16 total) for post hoc Pax6 and Nestin immunolabeling. Three samples were excluded because of cell loss during processing, and one additional sample was excluded because of misalignment between live-imaging and post hoc imaging fields. After quality-control filtering, the final dataset comprised 12 samples. Confocal images were acquired using a Zeiss LSM 880 spectral confocal microscope equipped with an LD LCI Plan-Apochromat 25x/0.8 objective. To align calcium imaging and confocal datasets, images were rescaled using a 0.7x correction factor to compensate for differences in magnification between acquisition systems. Image orientation was standardized prior to analysis, and ln2PMMA samples were rotated and/or mirrored when necessary to align groove orientation across datasets (**Fig. S5A**).

### Calcium Imaging analysis

Calcium imaging data were analyzed using a custom MATLAB-based algorithm to extract nuclear calcium signals from defined regions of interest (ROIs). ROIs of identical size were manually positioned over each nucleus to enable direct comparison of fluorescence intensities across cells. To correlate calcium dynamics with nuclear morphology and marker expression, snapshot images acquired during live imaging were aligned with corresponding post hoc immunolabeled images using ImageJ (representative example in **Fig. S5B)**. A total of 529 nuclei were successfully matched between calcium imaging and post hoc immunofluorescence datasets.

Background fluorescence was subtracted from each nucleus at every time point. Background values were calculated as the mean fluorescence intensity of 10 ROIs positioned in cell-free regions of the field of view. Background traces were inspected to identify potential imaging artifacts (**Fig. S5C**).

Calcium traces were smoothed using a Savitzky-Golay filter (polynomial order = 2, window size = 6) implemented in OriginPro v9 (OriginLab) (**Fig. S5D**). To minimize acquisition-related artifacts, the first 50 frames were excluded from analysis. Baseline fluorescence (F₀) was defined as the mean fluorescence intensity during the subsequent 50-frame baseline period recorded under calcium-free conditions:

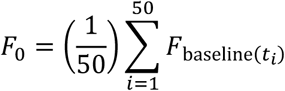

Relative fluorescence changes were calculated as:

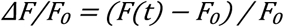

For visualization and quantitative analysis, values were expressed as percentage changes:

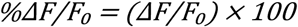

Unless otherwise indicated, all analyses were performed at the single-cell level. Parameters extracted from each nucleus included basal Ca²⁺ levels, transient frequency, and ΔF/F₀ amplitude. Nuclei were classified as active if they exhibited at least one detectable spontaneous Ca²⁺ transient during the recording period, whereas nuclei lacking detectable transients were classified as silent. Ca²⁺ transients were identified when fluorescence changes exceeded a predefined threshold of 3% above baseline.

### Immunofluorescence

Immunofluorescence was performed on cultured cells and tissue sections using sample-specific fixation and staining protocols.

Human radial glia (hRG) cultures were fixed in 4% paraformaldehyde (PFA) for 30 min at room temperature, rinsed in PBS, and blocked for 1 h in PBS containing 0.1% normal goat serum and 0.1% Triton X-100. Samples were incubated overnight at 4°C with primary antibodies diluted in blocking solution. Following washes in PBS, cultures were incubated with species-appropriate Alexa Fluor-conjugated secondary antibodies (Thermo Fisher Scientific) and counterstained with Hoechst 33342.

Human fetal forebrain tissue and mouse embryonic brains were fixed in 4% PFA, cryoprotected in 30% sucrose, embedded in OCT compound, and frozen. Cryosections (15 μm for human tissue and 40 μm for mouse tissue) were blocked and permeabilized in PBS containing normal serum and Triton X-100, incubated overnight at 4°C with primary antibodies, and subsequently incubated with Alexa Fluor-conjugated secondary antibodies and Hoechst 33342.

Mouse primary cortical cultures were fixed in 4% PFA in 0.1 M phosphate buffer for 20 min, permeabilized and blocked in PBS containing normal serum and Triton X-100, and incubated overnight at 4°C with primary antibodies. Following PBS washes, samples were incubated with Alexa Fluor-conjugated secondary antibodies and mounted in a Mowiol-based mounting medium.

hRG cultures were characterized by Sox2 and BLBP immunoreactivity, whereas neuronal populations were identified by TUJ1 and GABA expression. Additional marker combinations used in individual experiments are specified in the corresponding figure legends. Details of all primary antibodies, including source, catalog number, RRID, and working dilution, are provided in **Table S5**.

### Image processing and nuclear segmentation

Images were acquired using an LSM 880 spectral confocal microscope (Carl Zeiss, Germany) equipped with LD LCI Plan-Apochromat 25x/0.8 Imm Korr DIC M27 and Plan-Apochromat 40x/1.3 Oil DIC UV-IR M27 objectives. Image acquisition and z-stack reconstruction were performed using ZEN software (blue edition; Carl Zeiss). The number of optical sections was adjusted according to the experimental setup and markers analyzed. Identical acquisition and projection settings were applied across conditions for each marker. For each condition, six to nine images from two to four independent biological samples were analyzed. Images were acquired randomly across the substrate while avoiding regions of high cell confluence, where extensive cell-cell interactions may mask substrate-derived mechanical cues and promote astrocytic differentiation, as previously described^35^.

Nuclear segmentation was performed on either maximum– or ave-ensity z-projections, depending on the analysis. As both approaches yielded comparable morphometric measurements, they were considered equivalent for subsequent analyses (**Fig. S6A-B**). To optimize nuclear segmentation, different nuclear labeling strategies were evaluated. Lamin B1 immunostaining was compared with Hoechst staining, which increased nuclear detection under both control (16%) and ln2PMMA (18%) conditions without affecting measurements of nuclear area or eccentricity (**Fig. S6C-E**). Based on these results, Hoechst staining was selected as the standard nuclear label.

The segmentation pipeline combined global thresholding with additional refinement procedures, including a modified top-hat algorithm^77^, to delineate individual nuclei. From segmented nuclei, morphometric descriptors including area and eccentricity were extracted. Eccentricity (E) was defined as a dimensionless parameter ranging from 0 (circular) to 1 (maximally elongated), corresponding to the eccentricity of an equivalent fitted ellipse. The total number of nuclei per image was used as a proxy for local cell density. For orientation analysis, the angle between the major axis of each nucleus and the orientation of the ln2PMMA grooves was measured. In control substrates, angles were calculated relative to an arbitrary defined reference axis. Angles were transformed from a 0-360° range to a –90° to 90° range, where 0° corresponds to complete alignment, for analysis and visualization.

Quantitative image analysis was performed using a custom-designed MATLAB-based pipeline that enabled integration of marker expression measurements with nuclear morphometric features at the single-cell level. The analysis framework was designed to accommodate images acquired across different microscopy platforms through calibration based on acquisition parameters and user-defined inputs (**Fig. S7A, Table S4**). For each segmented nucleus, fluorescence intensities were extracted from nuclear and perinuclear regions of equal area. These measurements were used to calculate nuclear-to-cytoplasmic ratios for mechanosensitive signaling regulators (YAP/TAZ and β-catenin) and nuclear fluorescence intensities for Lamin proteins (**Fig. S7B-C**). To ensure accurate multi-channel quantification, segmentation masks were consistently applied across channels and manually validated to identify segmentation or alignment errors. For in vivo tissue sections and organoids, nuclear morphology was quantified manually using Fiji/ImageJ. Measurements were obtained from single optical planes or short z-projections (up to three sections). Cell density was estimated at the tissue-region level rather than on a per-image basis.

The numbers of biological samples, images, and nuclei analyzed for each experiment are reported in the corresponding figure legends.

### Modeling morphometric states associated with RG-like competence

A statistical modeling framework was implemented to quantitatively integrate morphometric parameters and assess their relationship with RG-associated states.

#### Model Overview

To quantitatively relate nuclear morphology to RG-associated marker expression, we implemented a generalized linear model (GLM) in R (R Core Team, Vienna, Austria). The model integrated morphometric and contextual descriptors extracted from confocal microscopy images of primary mouse cortical glial cultures maintained for 3DIV on either control or ln2PMMA substrates, together with measurements of RG-associated marker expression, as described above.

#### Predictor Variables

The model incorporated the following predictors:

- **Nuclear morphology**: nuclear area (A, µm²) and eccentricity (E, dimensionless), treated as continuous variables. To capture non-linear relationships, both variables were expanded using polynomial terms up to cubic order.
- **Cell density** (D): Defined as the number of nuclei per unit image area and used as a proxy for local cellular context.
- **Substrate condition** (S): categorical variable with two levels (control and ln2PMMA).

The dataset comprised 3,855 nuclei obtained from cortical glial cultures derived from 8-10 independent mouse litters across 15 independent experiments. Images were sampled across a range of confluence conditions to capture density-dependent effects.

#### Response Variable Definition

The response variable was defined using marker-informed categories employed exclusively for model training. For training purposes, nuclei were assigned to RG-associated or non-RG-associated groups based on validated combinations of lineage markers. RG-associated categories included Pax6+ nuclei (alone or in combination with Nestin, GFAP, or Sox2), as well as Sox2+/Nestin+ nuclei (n = 374). Non-RG-associated categories included marker-negative nuclei and lineage-committed populations (e.g., NG2+, Gsx2+, Pax6-/GFAP+) (n = 786). Cells with incomplete or ambiguous marker information (n = 2,695) were excluded to avoid introducing label uncertainty. Importantly, these marker-defined categories were used exclusively for supervised model training. The resulting model does not assign cellular identities but instead estimates the probability of occupying a continuous morphometric state associated with RG-like competence.

#### Model formulation

A binomial GLM with a logit link function was used to model the probability that a nucleus occupied a morphometric state associated with RG-like competence as a function of morphometric and contextual variables:

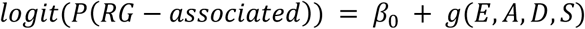

where *g(E, A, D, S)* represents polynomial expansions of nuclear eccentricity (*E*) and area (*A),* together with their interactions with substrate condition (*S*) and local cell density (*D*). Polynomial degrees and interaction terms were selected based on likelihood-based model comparison while maintaining biological interpretability. Importantly, inclusion of substrate-dependent interaction terms results in substrate-specific parameterizations within a unified modeling framework, such that morphometric predictors are weighted differently under control and ln2PMMA conditions (see **Table S3** for model details).

#### Model Selection and performance

Model selection followed an iterative likelihood-based reduction strategy guided by the Akaike Information Criterion (AIC), with the goal of identifying a parsimonious model while preserving explanatory power. Model reduction was primarily guided by AIC while retaining statistically significant terms and respecting hierarchical interaction structure.

Model performance was assessed using receiver operating characteristic (ROC) curves, and the area under the curve (AUC) was calculated separately for each substrate condition. Ninety-five percent confidence intervals were estimated by bootstrap resampling. Because substrate-dependent interaction terms were retained in the final model, distinct parameterizations were obtained for control and ln2PMMA conditions within a single unified framework. Full coefficient estimates and explicit model formulations are provided in **Table S3**.

#### Substrate-conditioned comparison and log-odds transformation

To directly compare how morphometric predictors relate to RG-associated probabilities (RGp) across substrates, we computed an odds ratio between conditions while holding morphometric variables constant. Taking the logarithm of this ratio yields the log-odds function:

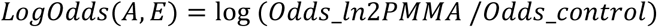

which depends only on nuclear area and eccentricity, as density-dependent terms cancel out during the comparison. This transformation provides a reduced representation of the model that captures substrate-dependent effects within morphometric space independently of the marker information used during training. It also enables direct visualization of substrate-associated differences through the LogOdds surface shown in **Fig. 5F**. The condition LogOdds = 0 defines a boundary, F(0), corresponding to equal probabilities under both substrate conditions. This boundary serves as a reference threshold for identifying regions of morphometric space associated with RG-like competence, as illustrated in **Fig. 5G**.

#### Validation strategy

Model validation was performed using two independent datasets derived from 3DIV glial cultures and processed using the same marker definitions as the training data. Estimated RG-associated fractions were compared with experimentally observed values to assess model consistency across biological replicates. Prior to model application, validation datasets were screened to ensure that predictor values fell within the range represented in the training data, and observations requiring extrapolation were excluded where appropriate.

Two complementary validation approaches were employed. First, fitted probabilities were analyzed as continuous distributions, and nuclei with RGp ≥ 0.5 were used as an operational threshold to estimate occupancy of the morphometric state identified by the model. Second, for datasets extending beyond the original training domain, (e.g., high-density cultures or *in vivo* samples), the LogOdds = 0 boundary was used to assess occupancy of morphometric space independently of probability calibration. Together, this dual strategy provides robust statistical validation while enabling application of the model across experimental contexts that differ in density, organization or sampling range. Accordingly, the model should be interpreted as a morphometric framework associated with RG-like competence rather than as a classifier of radial glial identity.

#### Publicly available single-cell transcriptomics data analysis

To characterize transcriptional dynamics associated with RG-astrocyte progression, we analyzed single-cell RNA-sequencing data from the mouse cerebral cortex generated by Di Bella et al.^43^ (GEO accession number GSE153162), focusing on genes related to mechanotransduction and calcium signaling. Data processing and analysis were performed using SCANPY^78^. Quality control filtering was applied based on the distributions of total UMI counts, number of detected genes, and mitochondrial transcript content. Cells with fewer than 500 or more than 50,000 UMIs, fewer than 800 or more than 7000 detected genes, or more than 15% mitochondrial reads were excluded. After filtering, 81,792 cells were retained from the initial 82,415 cells. An additional 2,431 cells identified as doublets using the Scrublet package^79^ were removed.

To focus the analysis on stages spanning the RG-to-astrocyte transition, cells from embryonic days E10-E12 were excluded, yielding a dataset of 67,961 cells. Expression values were normalized using the SCANPY normalize_total function to 10,000 counts per cell, followed by log-transformation. Dimensionality reduction was performed by principal component analysis (PCA), retaining the first 10 principal components. Cells were clustered using the Louvain community detection algorithm^80^ with a resolution parameter of 0.2. Clusters expressing RG-astrocyte lineage-associated markers (*Sox2, Pax6, Hes5*) and lacking expression of neuronal markers (*Neurod2, Tubb3, Neurod6, Meg3*) were selected for further analysis.

A second round of PCA was performed on the 14,469 selected cells, followed by clustering in the space of the first 11 principal components. Clusters expressing neuronal or oligodendrocyte lineage markers (*Olig1, Olig2, Pdgfra*) were excluded, with the exception of one cluster co-expressing *Olig1* together with mature astrocyte markers (*Aqp4, Slc1a2*; **Fig. S1**). Following this two-step selection process, 9,343 cells were retained as representative of the RG-astrocyte lineage. Expression of genes of interest was subsequently analyzed within this refined cell population. These analyses provided the basis for the marker-expression profiles and developmental expression patterns shown in **Fig. 2G-H**. Statistical analyses applied to both experimental and transcriptomic datasets are described below.

### Statistical analysis

Statistical analyses were performed using IBM SPSS Statistics (Version 26), GraphPad Prism (version 8.0), and R (R Core Team, 2021) within the RStudio environment (R Foundation for Statistical Computing, Vienna, Austria). Data distributions were assessed for normality, and statistical tests were selected accordingly. For comparisons between two groups, either Student’s t-test or the Mann-Whitney U test was used, depending on whether parametric assumptions were met. For comparisons involving three or more groups, one-way or two-way analysis of variance (ANOVA) followed by Tukey’s post hoc test was applied, or alternatively the Kruskal-Wallis test with appropriate multiple comparison procedures. Differences in categorical distributions were assessed using chi-square tests for analyses involving up to four groups. For contingency tables with more than four groups, a multiple regression-based approach adapted from previous work^81^ was employed. Statistical significance was defined as p < 0.05 with p < 0.01 considered highly significant. Model-specific statistical procedures are described in the corresponding methodological sections.

## RESOURCE AVAILABILITY

### Lead contact

Requests for further information and resources should be directed to and will be fulfilled by the lead contact, Soledad Alcántara.

### Materials availability

This study did not generate new unique reagents.

### Data and code availability

The primary scripts used to construct, apply, and visualizing the RGM model are available at https://github.com/PepPau/RGM-model-main-scripts.git.

Microscopy datasets generated during this study will be made available by the lead contact upon reasonable request.

## AUTHOR CONTRIBUTIONS

Conceptualization, S.A., J.P.S., J.A.O, and C.B.; Methodology, S.A., J.P.S.; Investigation, J.P.S. (mouse related experiments) and J.A.O. (human related experiments); Image acquisition and processing, J.P.S; Calcium imaging experiments, R.S. and J.P.S.; Software development for image and calcium analyses, C.B.; Data processing, J.P.S.; Data analysis, J.P.S. and S.A.; Statistical model generation, J.A. and J.P.S.; Single-cell transcriptomics meta-analysis, L.G.; Supervision, S.A., J.A.O. and C.B.; Writing – original draft, S.A. and J.P.S.; Writing – review and editing, all authors. All authors read and approved the final manuscript.

## ACKNOWLEDGMENTS

We thank the Biologia-Bellvitge Unit of the Scientific and Technological Centers (CCiTUB), Universitat de Barcelona, and Benjamin Torrejón for technical support and advice on confocal microscopy. We also thank Marta Cuenca and Cristina Martinez for their technical assistance in preparing human NPC cultures on control and ln2PMMA substrates. Finally, we thank Pau Gorostiza for hosting and supporting the calcium imaging experiments performed in his laboratory.

## SUPPLEMENTAL INFORMATION

Document S1. Containing Figures S1-S7 and tables S1-S5.

