## Supplementary figures and tables for "Mechanical cues stabilize a conserved morphometric state associated with radial glia-like competence across systems"

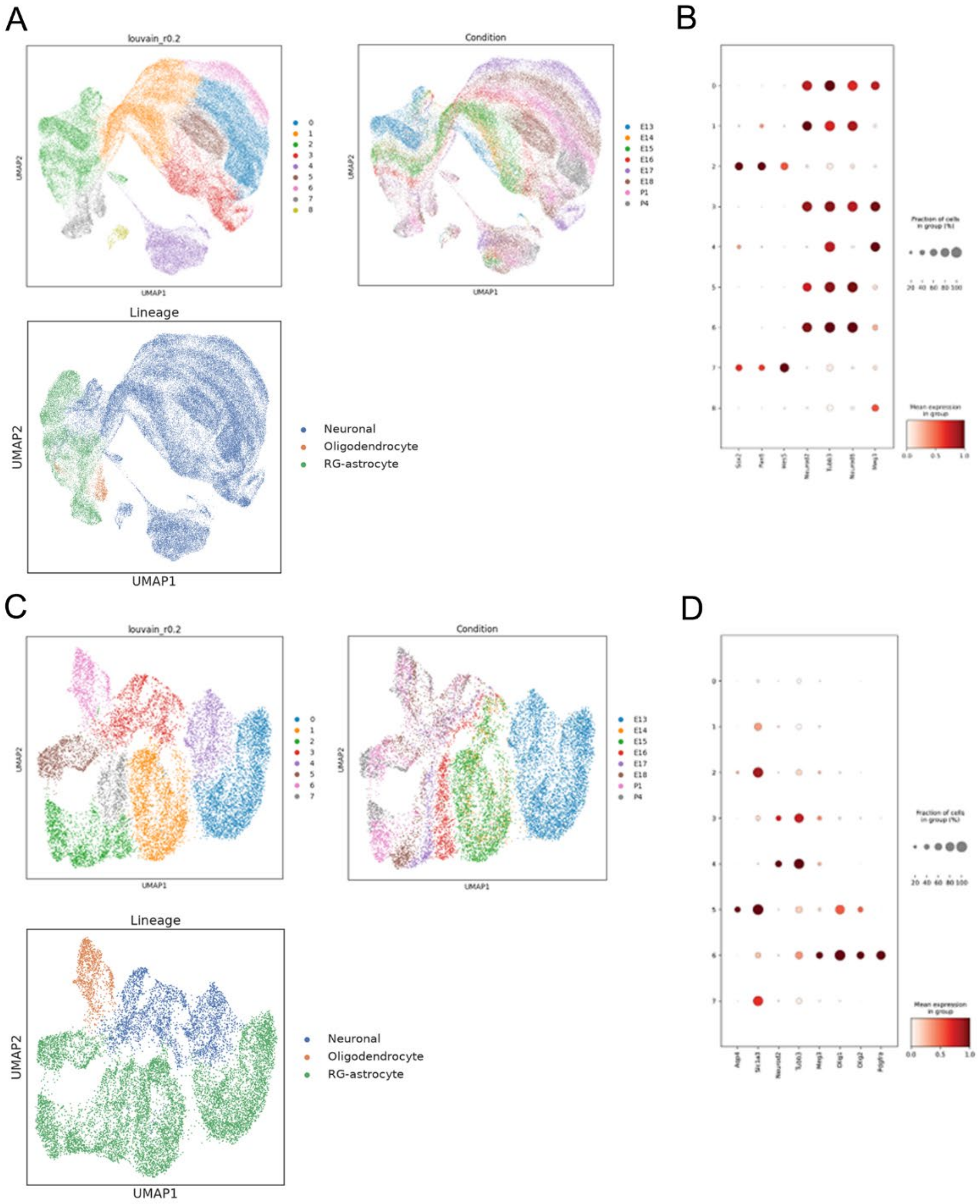

**Figure S1. Selection of RG-astrocyte lineage-associated cells for meta-analysis.**

**(A)** Uniform Manifold Approximation and Projection (UMAP) representation of nuclei from the dataset of Di Bella *et al*. (2021), colored by cluster (top left), developmental stage (top right), and expression of astrocyte, neuronal and oligodendrocyte marker genes (bottom).
**(B)** Expression of RG-astrocyte lineage-associated and neuronal marker genes across the clusters shown in (**A**). Based on these markers, clusters 2 and 7 were selected for further analysis.
**(C)** UMAP representation of the subset of cells selected on the basis of RG-astrocyte lineage-associated marker expression, colored by cluster (top left), developmental stage (top right), and expression of astrocyte, neuronal and oligodendrocyte marker genes (bottom).
**(D)** Expression of astrocyte, neuronal and oligodendrocyte marker genes across the clusters shown in (**C**). Based on these markers, clusters 3, 4, and 6 were excluded from further analysis.

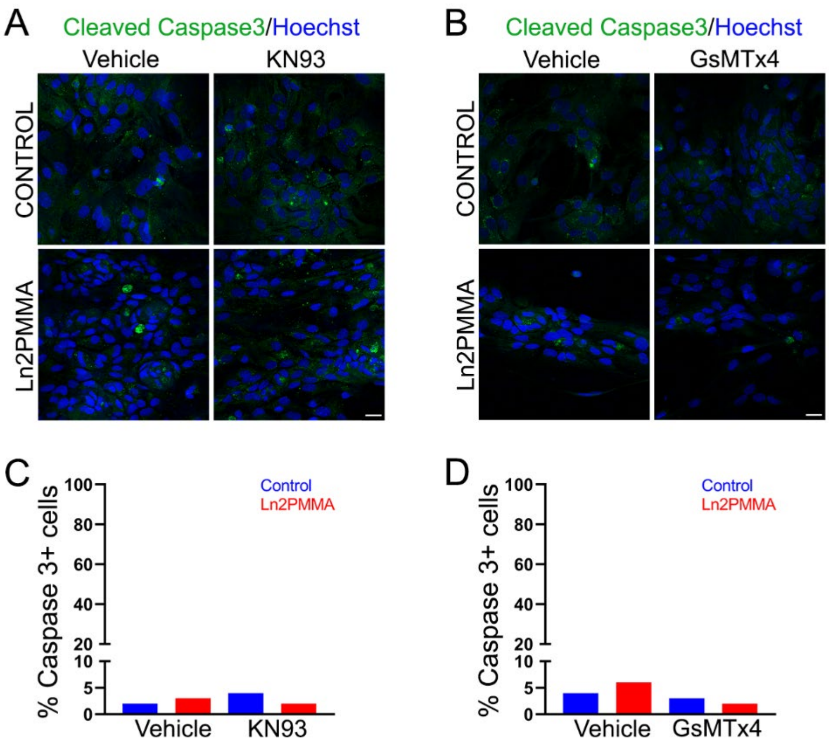

**Figure S2. KN93 and GsMTx4 treatments do not induce additional apoptosis in glial cells.**

**(A, B)** Representative confocal images of glial cells cultured on control and ln2PMMA substrates and immunostained for cleaved caspase-3, with nuclei counterstained with Hoechst 33342 (blue). Cells were treated with **(A)** vehicle (DMSO) or the CaMKII inhibitor KN93 (5 μM), and **(B)** vehicle (PBS) or the MSCs inhibitor GsMTx4 (1.22 μM).
**(C, D)** Quantification of the percentage of apoptotic cells (cleaved caspase-3-positive) on control and ln2PMMA substrates treated with **(C)** vehicle or KN93 and **(D)** vehicle or GsMTx4. Statistical significance was assessed using a chi-square test.
Scale bars = 20 μm.

**Table S1. Quantification of nuclear area and eccentricity of glial cells cultured for 3 DIV on control and ln2PMMA substrates and treated with KN93 or GsMTx-4.**

Cells were treated with KN93, GsMTx-4, or the corresponding vehicle controls (DMSO or PBS, respectively), and nuclear area and eccentricity were quantified. Statistical significance was assessed using Student's t-test: * p < 0.05, ** p < 0.01 compared with vehicle-treated cells on the same substrate.

| | | | Area | | | Eccentricity | | | |
|---|---|---|---|---|---|---|---|---|---|
| | | | **Mean** | **SD** | **p-value** | **Mean** | **SD** | **p-value** | **N** |
| **Nestin+ Pax6- cells** | **Control** | DMSO | 106.65 | 38.50 | | 0.75 | 0.13 | | 358 |
| | | KN93 | 97.27 | 37.58 | | 0.77 | 0.13 | | 317 |
| | | PBS | 99.35 | 42.94 | | 0.75 | 0.12 | | 241 |
| | | GsMTx-4 | 100.66 | 34.08 | | 0.76 | 0.11 | | 407 |
| | **Ln2PMMA** | DMSO | 86.34 | 29.94 | | 0.81 | 0.13 | | 324 |
| | | KN93 | 76.38 | 28.52 | ** | 0.81 | 0.12 | | 374 |
| | | PBS | 84.58 | 25.88 | | 0.81 | 0.12 | | 205 |
| | | GsMTx-4 | 91.79 | 29.25 | | 0.82 | 0.10 | | 347 |
| **Pax6+ cells** | **Control** | DMSO | 100.79 | 29.20 | | 0.75 | 0.13 | | 165 |
| | | KN93 | 97.11 | 38.44 | | 0.76 | 0.13 | | 173 |
| | | PBS | 99.69 | 39.13 | | 0.73 | 0.15 | | 66 |
| | | GsMTx-4 | 104.25 | 31.69 | | 0.75 | 0.13 | | 120 |
| | **Ln2PMMA** | DMSO | 83.18 | 25.72 | | 0.83 | 0.10 | | 227 |
| | | KN93 | 74.42 | 23.64 | | 0.83 | 0.12 | | 202 |
| | | PBS | 77.48 | 20.99 | | 0.85 | 0.12 | | 142 |
| | | GsMTx-4 | 90.37 | 26.92 | ** | 0.87 | 0.11 | | 128 |
| **Nestin- Pax6- cells** | **Control** | DMSO | 70.89 | 45.10 | | 0.67 | 0.15 | | 137 |
| | | KN93 | 75.21 | 41.59 | | 0.70 | 0.13 | | 112 |
| | | PBS | 68.12 | 42.71 | | 0.72 | 0.16 | | 155 |
| | | GsMTx-4 | 64.45 | 43.95 | | 0.70 | 0.14 | | 83 |
| | **Ln2PMMA** | DMSO | 64.34 | 32.72 | | 0.73 | 0.17 | | 128 |
| | | KN93 | 73.30 | 32.42 | | 0.76 | 0.14 | | 126 |
| | | PBS | 52.43 | 25.50 | | 0.72 | 0.15 | | 157 |
| | | GsMTx-4 | 57.03 | 25.90 | * | 0.73 | 0.17 | | 87 |

**Table S2. Quantification of nuclear area and eccentricity in marker-defined glial cell populations cultured for 3 DIV on control and In2PMMA substrates.**

Nuclear area and eccentricity were measured for each marker-defined cell population on the indicated substrates. Statistical significance between substrates for the same cell type was determined using Student's t-test. ** p < 0.01. Cell numbers correspond to marker-positive populations.

| | Cell type | Marker | Nº of cells (Control, In2PMMA) | Control Mean ± SD | In2PMMA Mean ± SD |
|---|---|---|---|---|---|
| AREA | NSC | Sox2 | 425 (236, 189) | 83 ± 39,4 | 86 ± 31,8 |
| | RG | Pax6 | 243 (91, 152) | 88 ± 30,8 | 88 ± 25,4 |
| | Tripotential IP | Gsx2 | 38 (15, 23) | 62 ± 17,3 | 56 ± 16,7 |
| | Astrocytes | GFAP | 362 (255, 107) | 95 ± 34,9 | 73 ± 25,2 ** |
| | Immature astrocytes/RG | Nestin | 1070 (527, 543) | 105 ± 44,9 | 88 ± 29,6 ** |
| | Oligodendrocyte IP (OPC) | NG2 | 68 (45, 23) | 59 ± 23,8 | 48 ± 28,1 |
| ECCENTRICITY | NSC | Sox2 | 425 (236, 189) | 0,75 ± 0,13 | 0,84 ± 0,12 ** |
| | RG | Pax6 | 243 (91, 152) | 0,78 ± 0,12 | 0,88 ± 0,10 ** |
| | Tripotential IP | Gsx2 | 38 (15, 23) | 0,65 ± 0,16 | 0,59 ± 0,15 |
| | Astrocytes | GFAP | 362 (255, 107) | 0,76 ± 0,12 | 0,85 ± 0,12 ** |
| | Immature astrocytes/RG | Nestin | 1070 (527, 543) | 0,76 ± 0,13 | 0,86 ± 0,10 ** |
| | Oligodendrocyte IP (OPC) | NG2 | 68 (45, 23) | 0,76 ± 0,14 | 0,82 ± 0,11 |

**Table S3. Summary of model performance, significant terms and equations.**

The top rows report the Akaike Information Criterion (AIC) value for the general model and the area under the curve (AUC) values stratified by substrate, including the lower and upper bounds of the 95% confidence intervals (CIs). The middle rows show the estimated coefficients, standard errors, z-values, and p-values for the selected model terms, grouped according to their substrate dependence. The bottom rows present the model equations. A, area; E, eccentricity; D, cell density.

| **Model** | **AIC** | **AUC** | **Lower 95% CI** | **Upper 95% CI** |
|---|---|---|---|---|
| General | 1300.325 | 0.73 | 0.7 | 0.76 |
| Control | | 0.67 | 0.62 | 0.71 |
| Ln2PMMA | | 0.77 | 0.73 | 0.81 |
| **Term** | **Estimate** | **Std. Error** | **z-value** | **p-value** |
| (Intercept) | -5.82E+03 | 7.62E+02 | -7.634 | 0 |
| **Dependent on substrates** | | | | |
| Ln2PMMA | -9.45E+02 | 5.66E+02 | -1.669 | 0.095 |
| Control * $E^3$ | -2.31E+02 | 8.84E+02 | -0.261 | 0.794 |
| Ln2PMMA * $E^3$ | 2.57E+03 | 7.32E+02 | 3.509 | 0 |
| Ln2PMMA * A | -7.33E+00 | 4.74E+00 | -1.546 | 0.122 |
| **Independent on substrates** | | | | |
| A | 1.51E+02 | 2.58E+01 | 5.829 | 0 |
| $A^2$ | -1.05E+00 | 2.22E-01 | -4.755 | 0 |
| $A^3$ | 1.81E-03 | 5.49E-04 | 3.296 | 0.001 |
| $E^2$ * D | 5.15E+00 | 2.49E+00 | 2.068 | 0.039 |
| $E^3$ * $D^2$ | -2.75E-03 | 1.50E-03 | -1.834 | 0.067 |
| A * D | -1.46E-01 | 4.13E-02 | -3.522 | 0 |
| A * $D^2$ | 8.80E-05 | 2.37E-05 | 3.722 | 0 |
| $A^2$ * D | 8.91E-04 | 2.78E-04 | 3.2 | 0.001 |
| $A^2$ * $D^2$ | -6.22E-07 | 1.84E-07 | -3.384 | 0.001 |
| **Model equations** | | | | |
| **Eq. control** | $\mathrm{logit}(\,P(y=1 \mid (X, \mathrm{Control}))) = \beta_0 + \beta_A A + \beta_{A^2} A^2 + \beta_{A^3} A^3 + \beta_{E^3 C} E^3 + \beta_{E^2 * D} E^2 * D + \beta_{E * D^2} E * D^2 + \beta_{A * D} A * D + \beta_{A * D^2} A * D^2 + \beta_{A^2 * D} A^2 * D + \beta_{A^2 * D^2} A^2 * D^2$ | | | |
| **Eq. ln2PMMA** | $\mathrm{logit}(\,P(y=1 \mid (X, \mathrm{ln2PMMA}))) = \beta_0 + \beta_{0Ln2} + \beta_A A + \beta_{A^2} A^2 + \beta_{A^3} A^3 + \beta_{E^3 Ln2} E^3 + \beta_{ALn2} A + \beta_{E^2 * D} E^2 * D + \beta_{E * D^2} E * D^2 + \beta_{A * D} A * D + \beta_{A * D^2} A * D^2 + \beta_{A^2 * D} A^2 * D + \beta_{A^2 * D^2} A^2 * D^2$ | | | |
| **Eq. LogOdds** | $Log(Odds\ ratio) = \beta_{0Ln2} + \beta_{E^3 Ln2} E^3 + \beta_{ALn2} A - \beta_{E^3 C} E^3$ | | | |

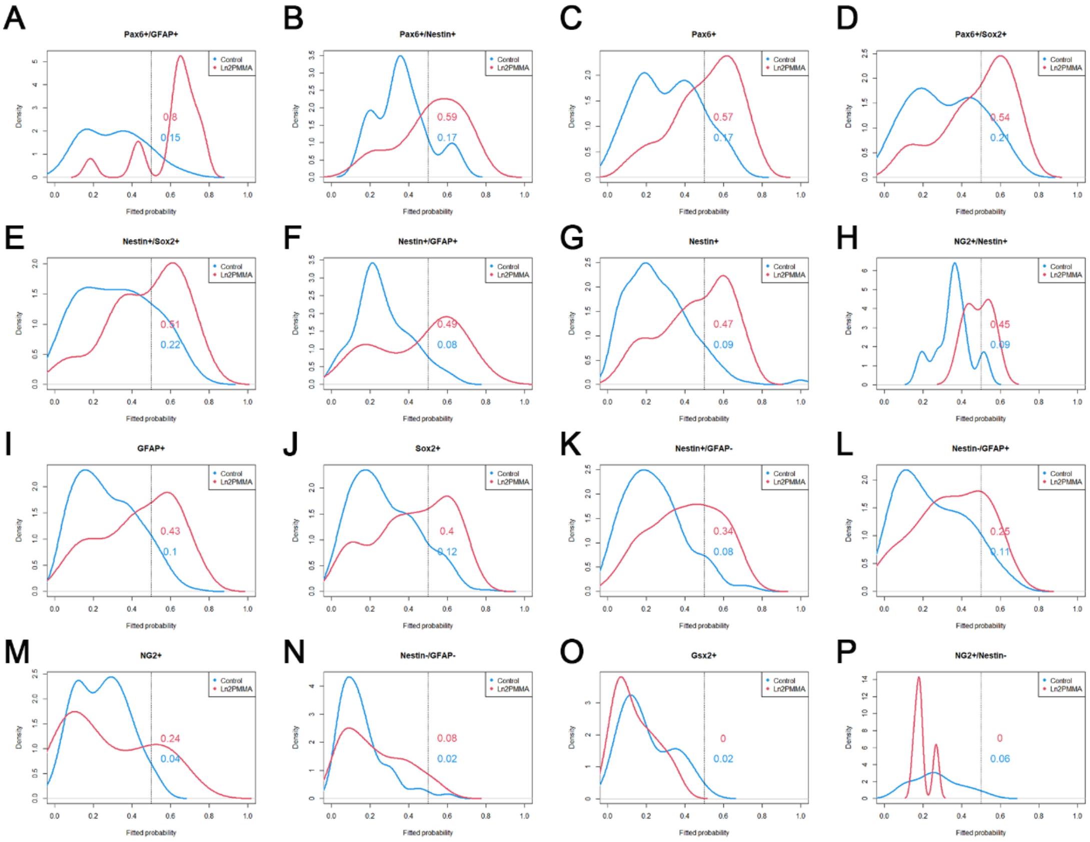

**Figure S3. Application of the RGM model to distinct glial subpopulations defined by lineage marker expression.**

Density plots show the RG-associated probabilities predicted by the RGM model for cells cultured on control (blue) or ln2PMMA (red) substrates using the dataset employed for model training. Subplots are ordered from left to right and top to bottom according to decreasing percentages of cells with RG-associated probabilities above the threshold (RGp = 0.5). Dashed lines indicate the RGp = 0.5 threshold. Numbers indicate the proportion of cells with RG-associated probabilities above the RGp = 0.5 threshold for each subpopulation.

**(A)** Pax6+/GFAP+ (n = 42), **(B)** Pax6+/Nestin+ (n = 50), **(C)** Pax6+ (n = 305), **(D)** Pax6+/Sox2+ (n = 103), **(E)** Nestin+/Sox2+ (n = 69), **(F)** Nestin+/GFAP+ (n = 68), **(G)** Nestin+ (n = 1,070), **(H)** NG2+/Nestin+ (n = 16), **(I)** GFAP+ (n = 362), **(J)** Sox2+ (n = 425), **(K)** Nestin+/GFAP- (n = 138), **(L)** Nestin-/GFAP+ (n = 84), **(M)** NG2+ (n = 68), **(N)** Nestin-GFAP- (n = 152), **(O)** NG2+/Nestin- (n = 12), **(P)** Gsx2+ (n = 38).

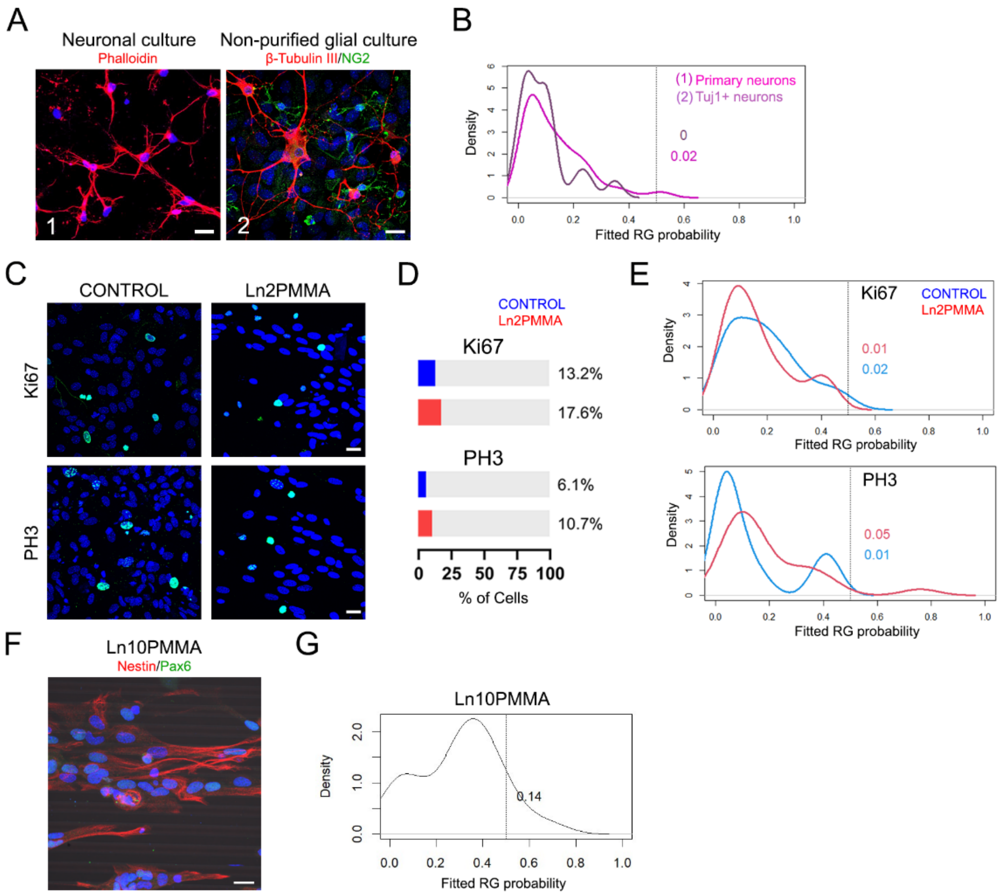

**Figure S4. Application of the RGM model to neuronal and mitotic subpopulations defined by marker expression and to glial cells grown on a distinct topography.**

**(A)** Representative images of neuronal cultures obtained from (1) embryonic day 16 cerebral cortices cultured on poly-D-Lysine coated borosilicate glass coverslips for 3DIV and stained with phalloidin and To-Pro-3 to visualize nuclei, and (2) βIII-tubulin-positive neurons in non-purified heterogeneous glial cultures at 7DIV, with nuclei counterstained with Hoechst 33342 (n = 18). Images were acquired using a Leica TCS-SL Spectral confocal microscope for condition (1) (n = 126) and a Zeiss LSM880 confocal microscope with a 40x objective for condition (2).

**(B)** Density plot showing the distributions of RG-associated probabilities predicted by the RGM model for primary neuronal cultures corresponding to conditions (1) and (2). Dashed line indicates the RGp = 0.5 threshold, and numbers indicate the proportion of cells with RG-associated probabilities above this threshold.

(**C**) Glial cultures grown on control and ln2PMMA substrates and immunostained for the proliferation markers Ki67 and phospho-histone H3 (PH3), with nuclei counterstained with Hoechst 33342.

**(D)** Bar graphs showing the percentage of Ki67-positive (control n = 108; ln2PMMA n = 54) and PH3-positive cells (control n = 42, ln2PMMA n = 24). Statistical comparisons were performed using a chi-square test.

**(E)** Density plots showing the distributions of RG-associated probabilities predicted by the RGM model for Ki67- and PH3-positive cells cultured on control (blue) and ln2PMMA (red) substrates. Dashed lines indicate the RGp = 0.5 threshold, and numbers indicate the percentage of cells with RG-associated probabilities above this threshold.

**(F)** Representative confocal image of glial cultures grown on PMMA micropatterned with 10 µm-wide grooves (ln10PMMA) immunostained for Nestin (red) and Pax6 (green), with nuclei counterstained with Hoechst 33342.

**(G)** Density plot showing the distribution of RG-associated probabilities predicted by the RGM model for glial cells cultured on ln10PMMA (22 nuclei). Numbers indicate the proportion of cells with RG-associated probabilities above the RGp = 0.5 threshold.

Scale bars = 20 µm.

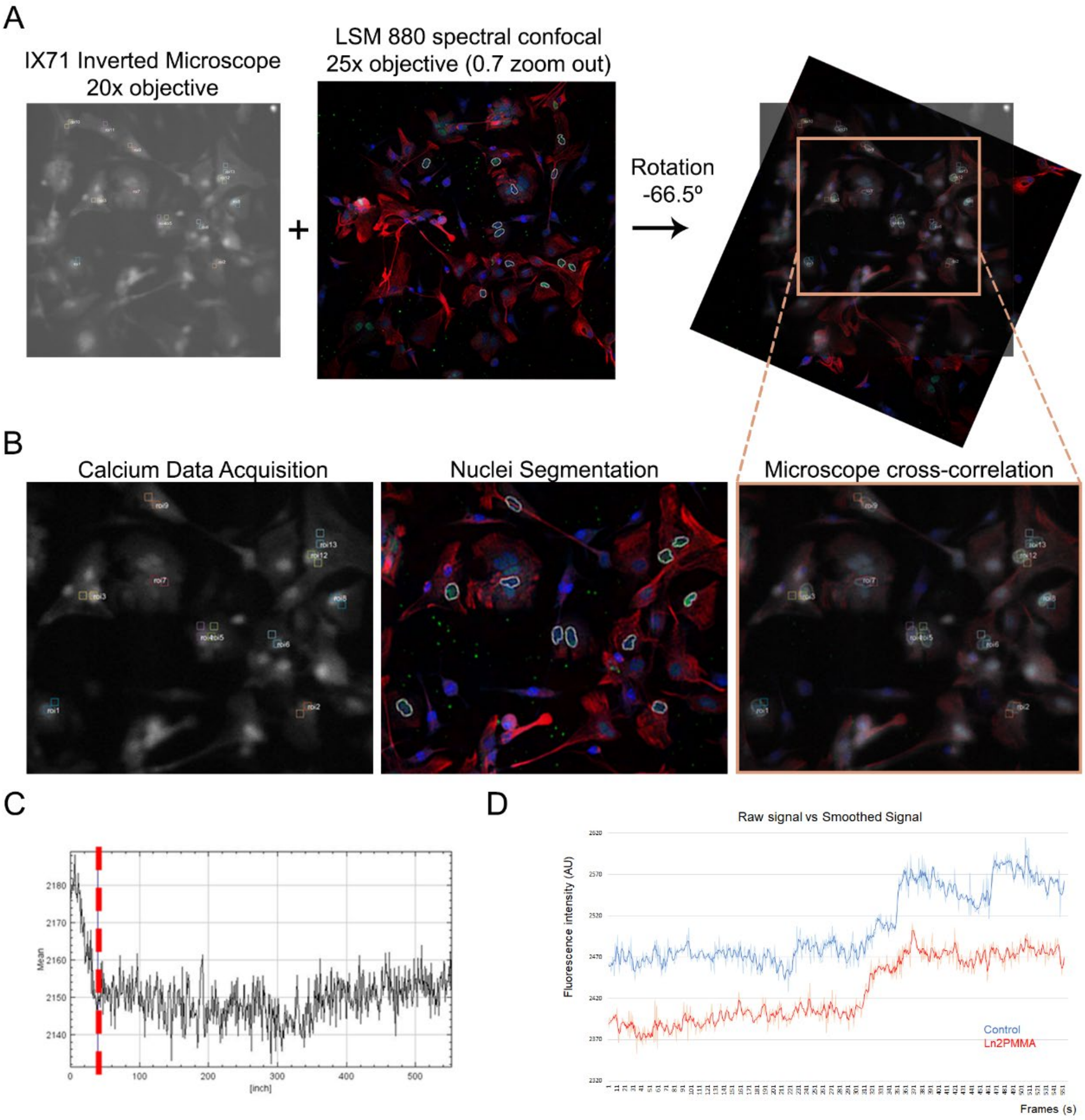

**Figure S5. Cross-correlation of live calcium imaging with post hoc immunolabeling, recording artifacts, and raw versus smoothed calcium signals.**

**(A)** Representative images acquired during live calcium imaging and the corresponding confocal images obtained after immunolabeling.
**(B)** Magnified view of the calcium imaging acquisition interface illustrating the correspondence between segmented nuclei and molecular marker expression (Nestin in red, Pax6 in green, and Hoechst in blue) used for nuclear identification. For each cell, two regions of interest (ROIs) of identical size and distance between them were defined; only the ROI located over the segmented nuclear region was included in the analysis.
**(C)** Representative calcium trace illustrating background fluorescence and initial recording artifacts. Data acquired before the red dashed line were excluded from subsequent analyses to eliminate acquisition-related artifacts.
**(D)** Representative calcium traces showing the raw signal and the corresponding smoothed signal (bold line) obtained using a Savitzky-Golay filter. Traces correspond to randomly selected cells cultured on control and ln2PMMA substrates.

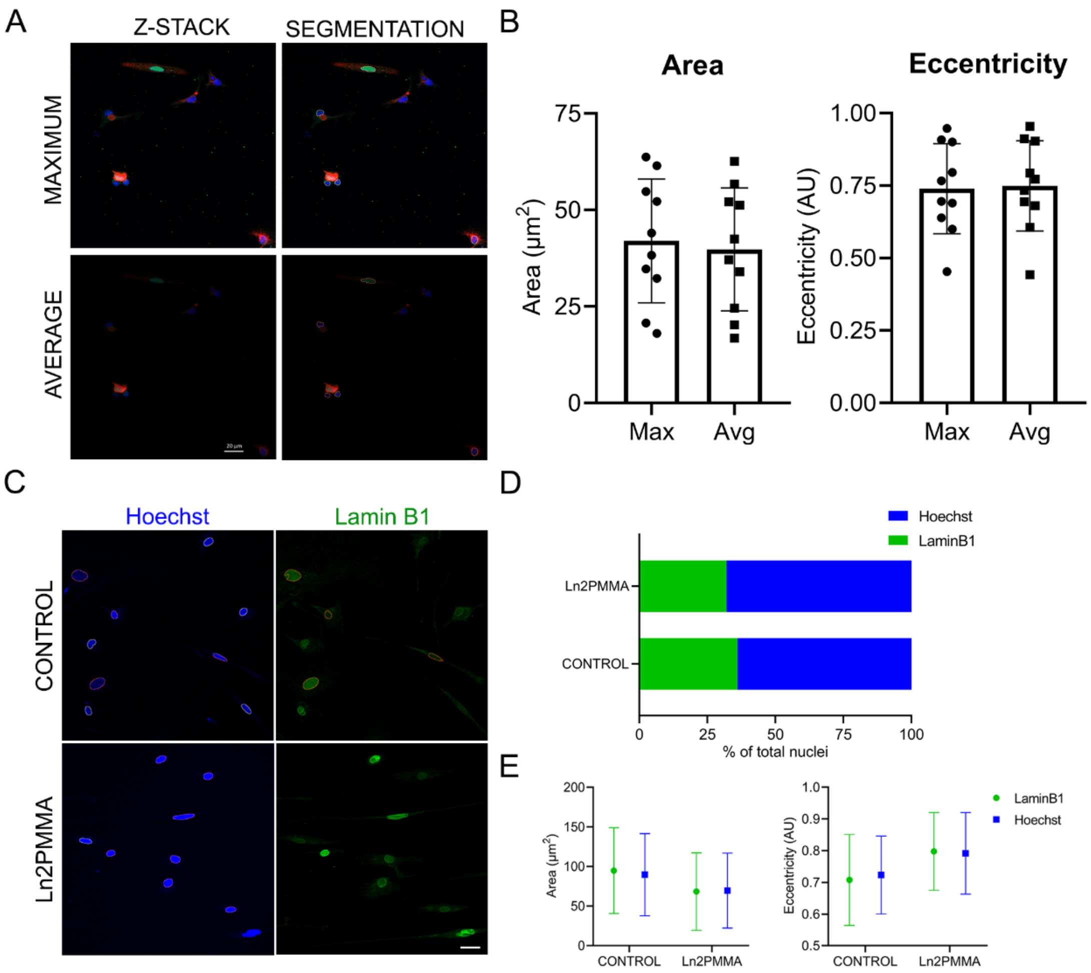

**Figure S6. Validation of nuclear segmentation robustness and labeling strategy.**

**(A)** Representative immunofluorescence images showing maximum- and average-intensity z-stack projections and their corresponding nuclear segmentations, demonstrating comparable segmentation performance between projection methods.

**(B)** Quantification of nuclear area and eccentricity derived from segmented nuclei using maximum- and average-intensity projections. Each point represents an individual nucleus. Statistical comparisons were performed using an unpaired Student's t-test (area, $p = 0.76$; eccentricity, $p = 0.89$).

**(C)** Representative immunofluorescence images of glial cultures maintained for 3 DIV on control and ln2PMMA substrates, with nuclei immunolabeled for Lamin B1 (green) and counterstained with Hoechst (blue).

**(D)** Percentage of successfully segmented nuclei obtained using Hoechst or Lamin B1 labeling. Under control conditions, 88 nuclei were analyzed (Hoechst, n = 61; Lamin B1, n = 47); under ln2PMMA conditions, 22 nuclei were analyzed (Hoechst, n = 17; Lamin B1, n = 13).

**(E)** Quantification of nuclear area and eccentricity (mean ± SD) obtained from Hoechst- and Lamin B1-labeled nuclei. Statistical comparisons were performed using an unpaired Student's t-test (control area $p = 0.71$; ln2PMMA area $p = 0.27$; control eccentricity $p = 0.54$; ln2PMMA eccentricity $p = 0.89$).

Scale bars = 20 μm.

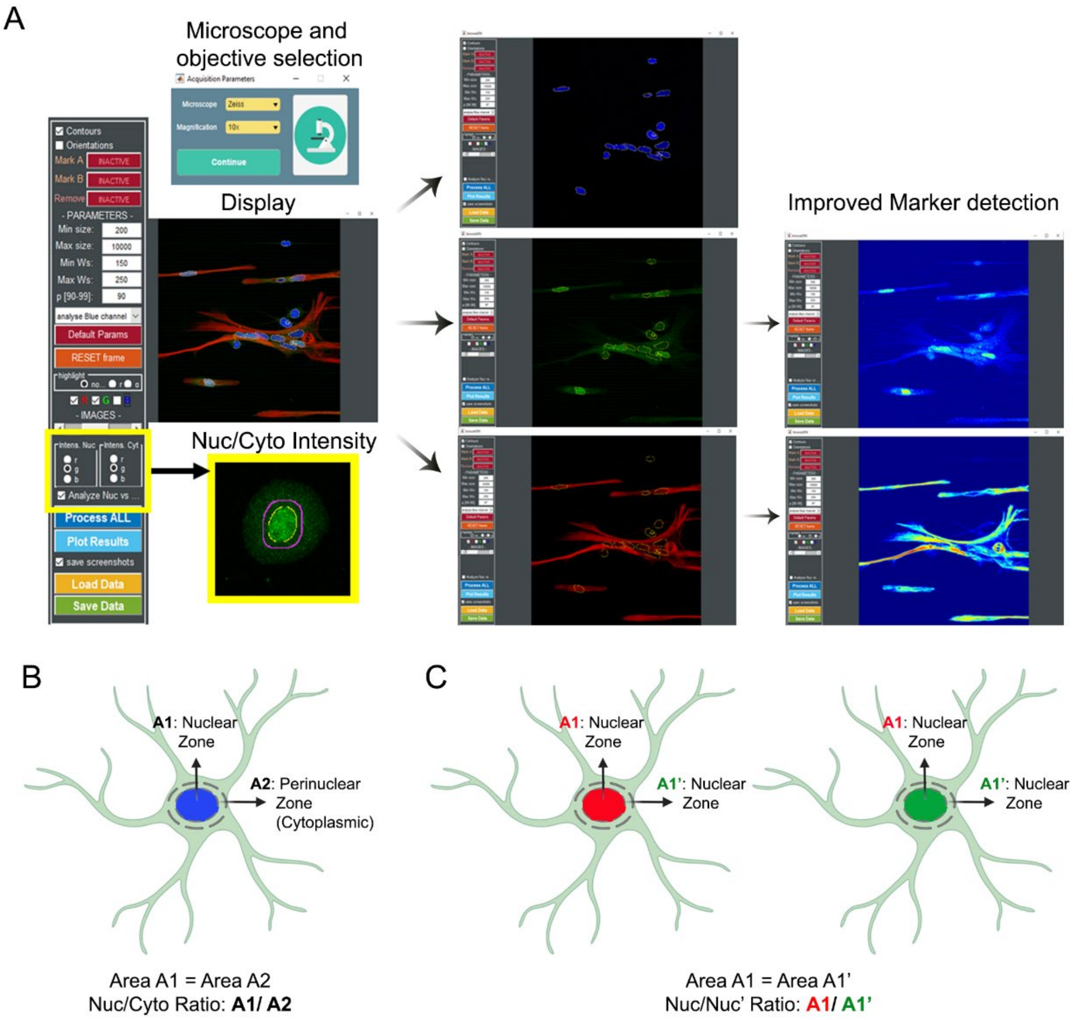

**Figure S7. Nuclear segmentation pipeline and schematic representation of nuclear and perinuclear fluorescence intensity extraction.**

**(A)** Overview of the in-house MATLAB application used for automated nuclear segmentation based on image acquisition settings and user-defined parameters. The interface allows visual inspection and manual correction of segmentation artifacts. The left panel contains controls for parameter adjustment and marker selection, including an optional module for extracting nuclear and perinuclear fluorescence intensity data (highlighted by a yellow-outlined box), whereas the right panel is dedicated to image visualization.
**(B)** Schematic representation of the workflow used to extract nuclear and perinuclear fluorescence intensities. Nuclear and perinuclear regions of identical size are defined independently and can be applied to any fluorescence channel.
**(C)** Schematic representation of the workflow used to extract nuclear fluorescence intensity measurements from two distinct imaging channels for the same segmented nuclei.

**Table S4. Objective-dependent conversion factors for microscopes used in this study.**

List of microscopes available in our facility and the corresponding objective-specific conversion factors used for image calibration and quantitative image analysis.

| Microscope | µm/pixel | Zoom |
|---|---|---|
| Nikon Eclipse 800 | 444.84/1360 | 20x |
| Zeiss Axiovert 200M | 161.4/1024 | 20x |
| Zeiss Apotome Axio Imager.M2 | 354.66/2056 | 20x |
| Leica TCS-SL (Confocal) | 375/1024 | 40x |
| Zeiss LSM 880 (Confocal) | 212.55/1024 | 40x |

**Table S5. Primary antibodies used in this study**

| Antibody | Host | Dilution | Supplier | Catalog No. | RRID | Antigen/marker |
|---|---|---|---|---|---|---|
| GFAP | Rabbit | 1:3000 | Dako (Agilent) | Z0334 | AB_10013382 | Mature astrocytes |
| NG2 | Rabbit | 1:600 | Millipore | AB5320 | AB_91789 | Oligodendrocyte progenitor cells |
| Pax6 | Rabbit | 1:500 | Abcam | ab195045 | AB_2750924 | Radial glia-associated transcription factor |
| Pax6 | Mouse | 1:500 | Santa Cruz Biotechnology | sc-81649 | AB_1127044 | Radial glia-associated transcription factor |
| Sox2 | Rabbit | 1:3000 | Abcam | ab97959 | AB_2341193 | Neural stem/progenitor cells |
| GSX2 | Rabbit | 1:500 | Invitrogen | PA5-40504 | AB_2605622 | Intermediate progenitors |
| Nestin | Mouse | 1:3500 | BD Biosciences | 556309 | AB_396354 | Radial glia and neural progenitors |
| Ki67 | Rabbit | 1:500 | Abcam | ab15580 | AB_443209 | Proliferating cells |
| Phospho-Histone H3 (PH3) | Rabbit | 1:500 | Millipore | 06-570 | AB_310177 | Mitotic cells |
| Lamin A/C | Mouse | 1:3000 | Thermo Fisher Scientific | 14-9847-82 | AB_2802197 | Nuclear envelope |
| Lamin B1 | Rabbit | 1:3000 | Abcam | ab16048 | AB_443298 | Nuclear envelope |
| YAP/TAZ | Rabbit | 1:1000 | Cell Signaling Technology | 8418 | AB_10950494 | Mechanotransduction effectors |
| β-Catenin | Rabbit | 1:3000 | Abcepta | AX10009 | n.d. | Canonical Wnt signaling |
| Sox2 | Goat | 1:1000 | Santa Cruz Biotechnology | sc-17320 (Y-17) | AB_2286684 | Neural stem/progenitor cells |
| BLBP | Rabbit | | Abcam | ab27171 | AB_869739 | Radial glia |
| GABA | Rabbit | | Sigma | A2052 | AB_477685 | GABAergic neurons |
| TUJ1 (βIII-tubulin) | Mouse | 1:10000 | Covance | MMS-435P | AB_2313773 | Neurons |

*Abbreviations: BLBP, brain lipid-binding protein; GFAP, glial fibrillary acidic protein; NG2, neuron-glial antigen 2; PH3, phospho-histone H3; TUJ1, class III β-tubulin. n.d., RRID not available.*
